# Characterization of Atypical Ebola Virus Disease in Ferrets

**DOI:** 10.64898/2026.01.20.700506

**Authors:** Wenguang Cao, Shihua He, Helene Schulz, Jordan Wight, Michael Chan, Karla Emeterio, Guodong Liu, Jonathan Audet, Kevin Tierney, Kimberly Azaransky, Kathy Frost, Lilianne Gee, Peter McQueen, Patrick Chong, Stephanie Booth, Garrett Westmacott, Erica Ollmann Saphire, Xiankun Zeng, Logan Banadyga

## Abstract

Ebola virus (EBOV) typically results in a severe—and often lethal—acute disease. However, increasing evidence suggests that EBOV can persist in certain immune-privileged tissues, which may then serve as reservoirs for the later reemergence of EBOV and disease recrudescence. Here, we report atypical EVD recrudescence in a ferret model inoculated with an otherwise lethal dose of EBOV and treated with low doses of a highly potent monoclonal antibody cocktail. Among 32 antibody-treated ferrets, 14 animals survived, while 8 succumbed to acute EVD within about 5-8 days. The remaining 10 animals succumbed to atypical EVD between 12 and 18 days post-infection (DPI) despite having shown no, or very minor, signs of illness during the acute phase of disease. While viremia disappeared by 14 DPI in most animals that succumbed to atypical EVD, it rebounded modestly just prior to death. Unlike animals that died of acute EVD, those that died of atypical EVD showed only a moderate systemic inflammatory response and few signs of organ dysfunction, in line with low levels of virus in the liver and spleen. Interestingly, however, ferrets that died of atypical EVD showed high levels of virus in the brain, consistent with increased markers of inflammation in the central nervous system and significant pathological changes, including a breakdown in the blood-brain barrier and severe meningoencephalitis. Not only does this study shed important light on the atypical and underappreciated manifestations of EVD, but it also establishes the ferret as a valuable model of EBOV persistence and recrudescence.

**AUTHOR SUMMARY:** Over the last several years, it has become increasingly apparent that Ebola virus is capable of persisting in survivors and, occasionally, reemerging to cause a recrudescent disease that is often distinct from the acute illness and characterized by neurological involvement. Since nearly all existing Ebola virus animal models are uniformly lethal, it has been difficult to understand the processes that lead to persistence and atypical manifestations of disease. In this study, we describe a late-onset, atypical Ebola virus disease in ferrets that was characterized by high levels of virus in the brain and significant markers of brain inflammation. Our comprehensive analysis of the pathogenic processes that contributed to this disease not only provides critical insight into Ebola virus persistence and recrudescence, but it also establishes the ferret as the first tractable model for investigating these atypical outcomes.

## INTRODUCTION

Filoviruses continue to pose a persistent threat to global public health and biosecurity. In particular, the orthoebolaviruses, which make up a genus within the family *Filoviridae,* are especially menacing. Ebola virus (EBOV), the prototypic orthoebolavirus, has been responsible for causing outbreaks of severe disease since at least 1976, when it was first discovered in what is now the Democratic Republic of the Congo (DRC) (1). For nearly 40 years, EBOV emerged sporadically throughout Central Africa, resulting in relatively small outbreaks with alarmingly high case fatality rates, exceeding 80% in some instances. In 2013, however, EBOV appeared for the first time in West Africa where it caused an epidemic that would last for over two and a half years, span multiple countries, infect nearly 30,000 people, and claim over 11,000 lives (2). Since then, EBOV has re-emerged regularly, mostly in Central Africa, to cause ten additional outbreaks, including the second largest filovirus outbreak on record, which sickened over 3,000 people and killed over 2,000 in the DRC from 2018 to 2020 (3).

Prior to the 2013-2016 West African EBOV epidemic, Ebola virus disease (EVD) was generally considered to be an acute disease. Despite well-documented long-term sequelae in some survivors, virus persistence (in semen) was rarely observed and EVD recrudescence or relapse was clinically unknown. However, the tens of thousands of EVD cases that resulted from the 2013-2016 epidemic not only resulted in an unprecedented number of deaths, but they also resulted in an equally unprecedented number of survivors—dramatically increasing the likelihood of observing otherwise very rare manifestations of EVD. Additionally, the advent of post-exposure treatments, including small molecule antivirals, monoclonal antibodies, and post-exposure vaccinations, likely allowed some patients to survive infections with viral loads that previously would have been fatal. Thus, in the aftermath of the large outbreaks of the past ∼15 years, EBOV is now understood to persist in a variety of immune-privileged sites, including the urogenital system, eyes, and the central nervous system (4). Virus persistence in the testes, and subsequent shedding in the semen, has been implicated in the sexual transmission of EBOV (5–14), while persistence in the eyes and central nervous system has been linked to recrudescence and organ-specific inflammatory diseases, such as uveitis and meningoencephalitis (15, 16). Unfortunately, the mechanisms that lead to persistence, followed by the pathophysiological changes that contribute to recrudescence, are not fully understood. While work in nonhuman primates (NHPs) has shed some light on these phenomena (17), the rarity and unpredictability of their presentation make it difficult to generate conclusive data.

Thankfully, the past several years of filovirus research have produced a number of new animal models, which are critical for better understanding virus pathogenesis and evaluating novel countermeasures (18). The ferret model, in particular, has been developed by multiple groups for a number of orthoebolaviruses, including EBOV (19, 20). Unlike the rodent models of orthoebolaviruses, which require either immunocompromised animals or host-adapted viruses, ferrets, like humans and NHPs, are susceptible to infection caused by wild type viruses (21). In the case of EBOV, disease in ferrets is rapid and severe, with animals typically succumbing within 5 or 6 days and displaying several of the hallmarks of EVD, including unchecked and systemic virus replication, a dysregulated immune response, and, eventually, multiorgan failure (21). Remarkably, EVD presentation in ferrets closely parallels EVD in NHP models, which display all the same hallmarks of infection but succumb to disease slightly later, typically on 7-8 days post infection, depending on the species (22). Despite the potential for ferrets as an intermediate animal model for orthoebolaviruses, relatively few studies have used these animals to assess countermeasure efficacy, particularly for EBOV. While the absence of many ferret-specific reagents has undoubtedly contributed to this lack of use, the fact that ferrets are highly sensitive to severe disease caused by EBOV infection likely also makes these animals a challenging model in which to conduct successful countermeasure evaluation.

We initially set out to benchmark the efficacy of a proven human monoclonal antibody (mAb) cocktail under a variety of treatment and infection conditions in ferrets, noting that the disease course is rapid, the half-lives of human mAbs in ferrets are short, and species mismatch may preclude Fc-mediated protection. We chose to evaluate an intranasal (IN) route of inoculation in addition to an intramuscular (IM) route, reasoning that inoculation of a mucous membrane may delay the onset of disease and increase the therapeutic window. We also chose multiple treatment times, ranging from very early treatment, on days 2 and 4 post-inoculation, to relatively late treatment times, on days 3 and 6.

The outcome of the experiments revealed a correlation between the circulating level of the mAb cocktail and survival, with higher levels of antibody linked to the absence of acute disease and survival and lower levels of antibody linked to the development of EVD and death. Interestingly, animals that exhibited intermediate levels of antibody did not develop acute disease but succumbed to a late-onset illness that bore little resemblance to typical EVD. Many of these animals exhibited signs consistent with neurological impairment immediately prior to death, with evidence of a pro-inflammatory immune response in the brain along with meningoencephalitis. Accordingly, virus levels in the brain were high at the time of death, despite much lower levels of viral RNA in the blood. Clinical chemistry, hematology, and cytokine parameters were moderately disturbed, showing both similarities and differences with the surviving animals as well as the animals that had died of acute EVD. Notably, the pattern of “atypical” EVD experienced by these ferrets bore several significant similarities to rare cases of atypical EVD in humans (15, 23, 24) and NHPs (25–28), which have also demonstrated disease relapse associated with encephalitis and meningoencephalitis.

Because mAbs reliably protect against death from EVD—in animal models and humans, alike—they may paradoxically create the opportunity to detect very rare manifestations of EBOV infection following recovery from acute disease. Given the unexpected findings presented here, these rare manifestations may be particularly easy to detect in ferrets, which are especially susceptible to severe disease caused by EBOV, difficult to effectively treat with human mAbs (owing to short half-lives and reduced effector function of human mAbs in ferrets), and relatively easy to study in large numbers (compared to NHPs). Our thorough characterization of atypical EVD in these animals therefore supports their use as a tractable and reproduceable model system in which to investigate the many mysteries that continue to surround EBOV persistence and recrudescence.

## RESULTS

### Treatment with human antibodies partially protected ferrets from EBOV

Two groups of ten ferrets each were inoculated with a target dose of 1000 TCID_50_ EBOV via either the IN or IM route, after which four animals in each group were treated with 30 mg/kg each of two human mAbs on days 2 and 5 post-infection (“early” treatment) or days 3 and 6 (“late” treatment) (Fig. S1A, B and Table S1). As controls, two ferrets from each inoculation group were treated with phosphate buffered saline (PBS) on 2 and 5 days post-infection (DPI). All four control animals developed severe disease and succumbed or were euthanized on day 5 (IM) and 6 (IN) post-infection, as expected (Fig. 1A). Three of the IM-inoculated and treated animals also developed severe disease, with one early-treated animal requiring euthanasia on 6 DPI and two late-treated animals requiring euthanasia on 7 and 8 DPI (Fig. 1A). None of the 13 animals remaining at 8 DPI had displayed any significant signs of disease up to that point, and all were gaining or maintaining their body weight (Fig. 1B). On 12 DPI, one animal (ID# 653M) in the IM-inoculated and late-treated group began to show signs of disease that progressed rapidly, with the animal meeting euthanasia criteria later the same day. Between 15 and 18 DPI, five more animals succumbed to disease that appeared suddenly and progressed rapidly: two animals that were IM-inoculated and treated early, one animal that was IM-inoculated and treated late, and two animals that were IN-inoculated and treated late. A total of seven animals survived until the end of the study on 29 DPI, including all four animals that were IN-inoculated and treated early. None of the survivors exhibited any pronounced signs of disease at any point after virus inoculation.

**Figure 1.**
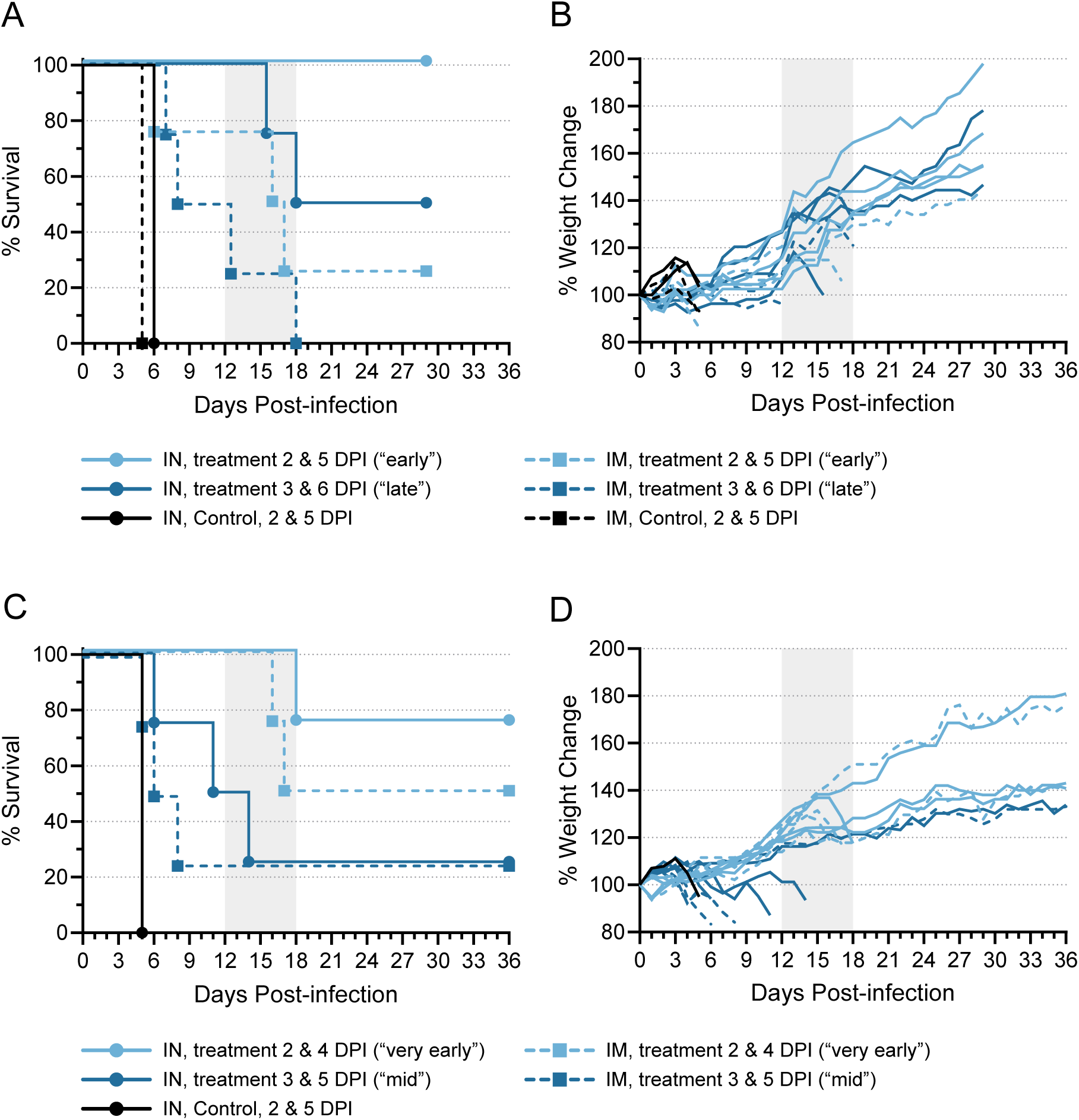
Human mAb provides limited protection from Ebola virus disease in ferrets. Ferrets were inoculated with EBOV variant Makona-C07 and treated with a human mAb cocktail in two independent experiments. In both the first (A, B) and second (C, D) experiment, ferrets were inoculated with a target dose of 1000 TCID_50_ EBOV via the intranasal (IN; n = 10/experiment) or intramuscular (IM; n = 10/experiment) routes. In the first experiment, 8 ferrets from each inoculation group were treated with 30 mg/kg each antibody delivered via the intraperitoneal (IP) route “early”, on 2 and 5 days post-infection (DPI), or “late”, on 3 and 6 DPI. Two ferrets from each inoculation group served as control animals and were treated with PBS only on 2 and 5 DPI. Animals were monitored for 29 days for survival (A) and weight change (B). In the second experiment, 8 ferrets from each inoculation group were treated with 30 mg/kg each antibody delivered via the IP route “very early”, on 2 and 4 DPI, or “mid”, on 3 and 5 DPI. One ferret from the IN group served as a control animal and was treated with PBS only on 2 and 4 DPI. Animals were monitored for 36 days for survival (C) and weight change (D). On all graphs, the vertical area shaded grey, from 12 to 18 DPI, represents the window in which atypical disease was observed.

We repeated the study and decreased the time between antibody doses from two days to one. Thus, animals were treated with 30 mg/kg of each human mAb on days 2 and 4 post-infection (“very early” treatment) or days 3 and 5 (“mid” treatment) (Fig. S1C, D and Table S1). A single control animal, which was IN-inoculated and treated with PBS on 2 and 4 DPI, succumbed to disease on 5 DPI (Fig. 1C). Four mid-treated animals—three that were IM-inoculated and one that was IN-inoculated—developed severe disease and were euthanized on 5, 6, and 8 DPI (Fig. 1C). Of the 12 remaining animals on 8 DPI, all but one appeared clinically normal, with no pronounced signs of disease observed up to that point and weights increasing (Fig. 1D). One animal (752M) from the IN-inoculated, mid-treated group showed signs of disease beginning around 4 DPI and remained ill until ultimately meeting euthanasia criteria on 11 DPI (Fig. 1C). As in the first experiment, four animals developed a rapid-onset disease late post-infection and were euthanized between 14 and 18 DPI: one IN- and two IM-inoculated animals treated very early and one IN-inoculated animal that received the mid treatment. A total of seven animals survived to study end, which was extended to 36 DPI for this study. Six of these animals showed no significant signs of disease throughout the experiment, while one animal (977F) in the IM-inoculated, mid-treated group became acutely ill between 3 and 6 DPI, but recovered completely by 7 DPI.

Of all the animals evaluated at different inoculation routes and doses, fewer than half survived (Table 1 and Table S1). However, we did note a difference in survival between animals inoculated via the IN route and animals inoculated via the IM route: 10 of the 16 (62.5%) IN-inoculated animals survived, whereas only 4 of the 16 (25%) IM-inoculated animals survived. Indeed, of the IN-inoculated animals treated early or very early, only one succumbed, suggesting that IN inoculation followed by early treatment may be the most effective way to demonstrate antibody efficacy in the ferret model.

**Table 1.**
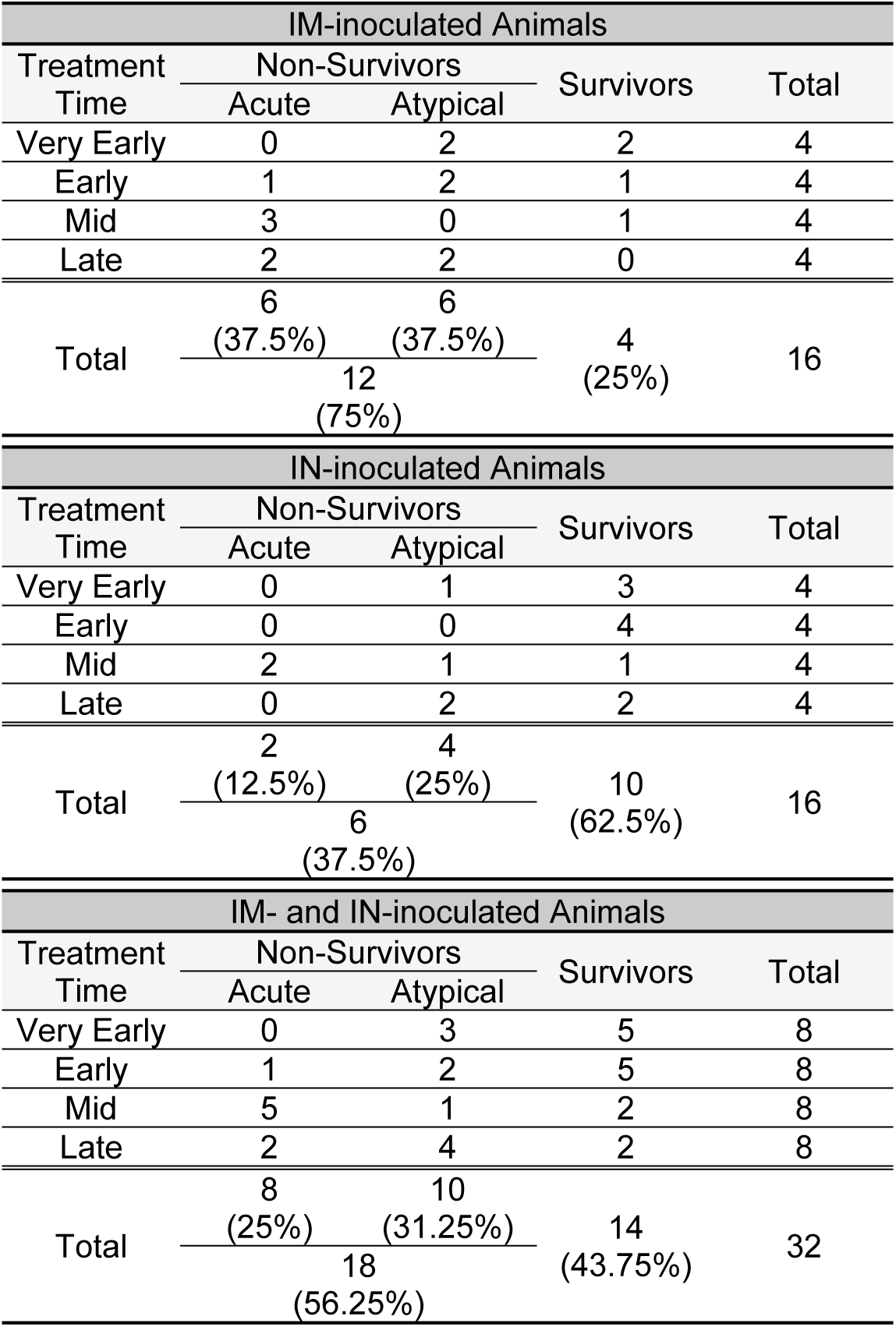
Outcomes in mAb-treated ferrets.

Among all 18 antibody-treated, non-surviving animals, only 7 (∼40%) succumbed to disease on day 8 or sooner, around the same time that the 5 untreated control animals died and consistent with previous results using the ferret model (21). One animal (752M) died on day 11, but considering that it first showed signs of disease on 4 DPI that persisted until euthanasia, this animal likely experienced a protracted form of acute disease. The remaining 10 of the 18 (∼60%) treated, non-surviving animals succumbed to disease between 12 and 18 DPI, with 7 animals succumbing as late as 16-18 DPI, despite showing no or very mild signs of disease early after inoculation. All 10 of these animals exhibited up to 15% weight loss over the day or two preceding euthanasia (Fig. 1B, D), and a few of them exhibited elevated temperatures, although none exhibited signs of disease requiring a clinical score until the day before—or even the day of—euthanasia. Disease in all of these animals appeared abruptly and progressed rapidly, with all animals exhibiting dramatically decreased activity. Several animals also exhibited signs consistent with neurological problems, including limited or no movement, rolling movement, lack of coordination, or the inability to right themselves. Based on clinical signs alone, it was clear that these animals did not succumb to disease that is typical for EBOV in ferrets. For clarity throughout the manuscript, we therefore refer to the disease experienced by all animals that died on 12 DPI or later as atypical EVD, and we use the term acute EVD to refer to the disease experienced by all animals that died on 11 DPI or earlier.

### Viremia rebounded with atypical EVD

In almost all animals, regardless of the route of inoculation or treatment strategy, viral RNA levels in the blood peaked on 4 or 5 DPI before decreasing substantially following mAb treatment (Fig. 2A). Viral RNA was reduced to very low or undetectable levels in most remaining animals by 10 DPI, and, by the end of the studies (29 or 36 DPI), viral RNA was no longer detectable in the blood of any of the survivors. Of the animals that experienced atypical EVD, viral RNA levels disappeared in eight of them by 14 DPI. The two animals (653M, 043F) that succumbed to atypical EVD earlier than the rest—on 12 and 14 DPI—showed drops in their viral RNA levels following mAb treatment, but the RNA was never cleared. After 14 DPI, however, viral RNA reappeared in the blood of all eight remaining atypical animals, coincident with their clinical deterioration.

**Figure 2.**
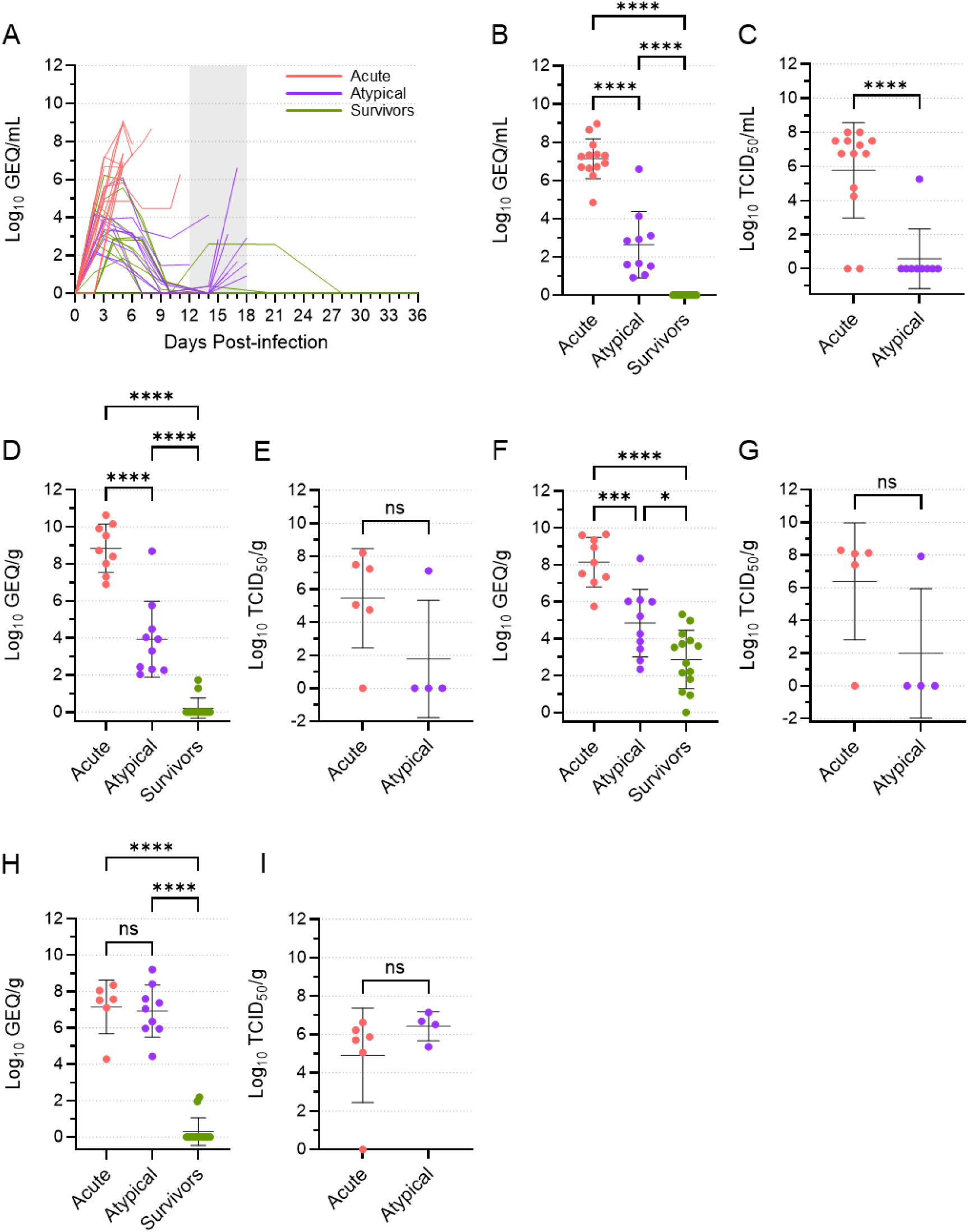
Levels of viral RNA and infectious virus differ among acute, atypical, and surviving ferrets. Viral RNA loads (genome equivalents [GEQ] per millilitre) in the blood were determined over time for the acute, atypical, and surviving ferrets (A). The vertical area shaded grey, from 12 to 18 DPI, represents the window in which atypical disease was observed. Viral RNA loads (in GEQ/ml or GEQ/g) at the terminal time points are depicted separately for the blood (B), liver (D), spleen (F), and brain (H) samples, with means and standard deviations indicated. Mean levels were compared using a one-way ANOVA with Tukey’s multiple comparison test. Infectious virus loads at the terminal time points are depicted in median tissue culture infectious dose (TCID_50_) per ml or g, with means and standard deviations indicated, for the blood (C), liver (E), spleen (G), and brain (I) samples. Mean levels were compared using a t test. ns, not significant; *, p≤0.05; ***, p≤0.001; ****, p≤0.0001.

To compare viral RNA levels among the three different disease groups, we grouped all data from the terminal time points of animals that experienced acute disease, animals that experienced atypical disease, and animals that survived. This analysis revealed that, although viral RNA levels in the atypical animals increased just prior to their death, the mean viral RNA load in the blood of these animals was still significantly lower than what was observed in the acute animals (Fig. 2B). A similar pattern was observed with respect to infectious virus found in the blood (Fig. 2C). The majority of acute animals had 6-8 log_10_ TCID_50_/ml of virus at the terminal time point, although infectious virus was not recovered from two animals. Conversely, infectious virus was only recovered from a single blood sample from an atypical animal (698M), with all other samples negative. Thus, animals were dying of atypical EVD with significantly lower levels of viremia than those dying of acute EVD, suggesting that other factors may be influencing disease course.

### Animals dying of atypical EVD had high levels of virus in the brain

We next examined the systemic spread of the virus using a limited set of liver, spleen, and brain samples taken from most ferrets at the time of euthanasia (Fig. 2D-I). Similar to what was observed in the blood, viral RNA levels were significantly higher in the liver and spleen from the acute animals than they were from the atypical animals, suggesting that infection of these organs was much more active in the animals that died of acute disease (Fig. 2D, F). This conclusion was supported by the isolation of infectious virus from 5 of 6 liver and spleen samples from acute animals, compared to only 1 of 4 liver and spleen samples from atypical animals (Fig. 2E, G), although the differences in mean levels of infectious virus were not statistically significant. Lower levels of viral RNA were detected in most spleen samples assessed from the surviving animals, but no infectious virus was detected in the 6 samples that were available for virus isolation, suggesting that the presence of viral RNA was not the result of a concurrent productive infection. Strikingly, the levels of viral RNA in brain samples from the acute and atypical animals were essentially equal and significantly higher than what was observed in the surviving animals, where most animals had no detectable viral RNA in the brain (Fig. 2H). These data were corroborated by the isolation of infectious virus from 5 of 6 brain samples from the acute animals and all four brain samples from the atypical animals (Fig. 2I), and they suggest that both acute and atypical animals experienced high levels of virus replication in the brain.

Virus genomes were sequenced from the brain samples of six acute animals and nine atypical animals, as well as from the liver samples of nine acute animals and ten atypical animals (Fig. S2). A few synonymous and non-synonymous mutations were identified in some of the samples, but almost all mutations were present at very low frequencies and none were present only in acute or atypical animals. A non-synonymous mutation in VP24 (E180V) was identified in many of the brain and liver samples, mostly from the acute animals; however, with the exception of three animals (one acute and two atypical), the frequency of this mutation was very low. Two GP mutations (T544I, D552N) and one VP30 mutation (t9267c) were present in the inoculum virus at relatively high frequences. The frequencies of these mutations increased slightly in the atypical animals compared to the acute animals, but the relevance of this increase is unclear.

### Severe meningoencephalitis, ventriculitis, and choroid plexitis in animals with atypical EVD

To further characterize the brain infection in animals with atypical EVD, we performed RNA *in situ* hybridization to detect EBOV genomic RNA in brain tissues. In the ferrets that died of acute EVD, EBOV RNA was limited in the endothelial vessels of the brain (Fig. 3A, B), which are primary targets of virus infection. In contrast, in animals that succumbed to atypical EVD, EBOV RNA and protein were highly abundant in the infiltrates of the meninges, choroid plexuses, ventricles, and their adjacent neuropils (Fig. 3C, D). No viral RNA was detected in the endothelial vessels of the brains in these animals. Histological analysis of brain tissues from the atypical animals demonstrated meningoencephalitis, ventriculitis, and choroid plexitis characterized by infiltrates of lymphocytes, plasma cells, and macrophages in the meninges, ventricles, and their adjacent cerebral perivascular spaces and in the choroid plexuses (Fig. 3E-H).

**Figure 3.**
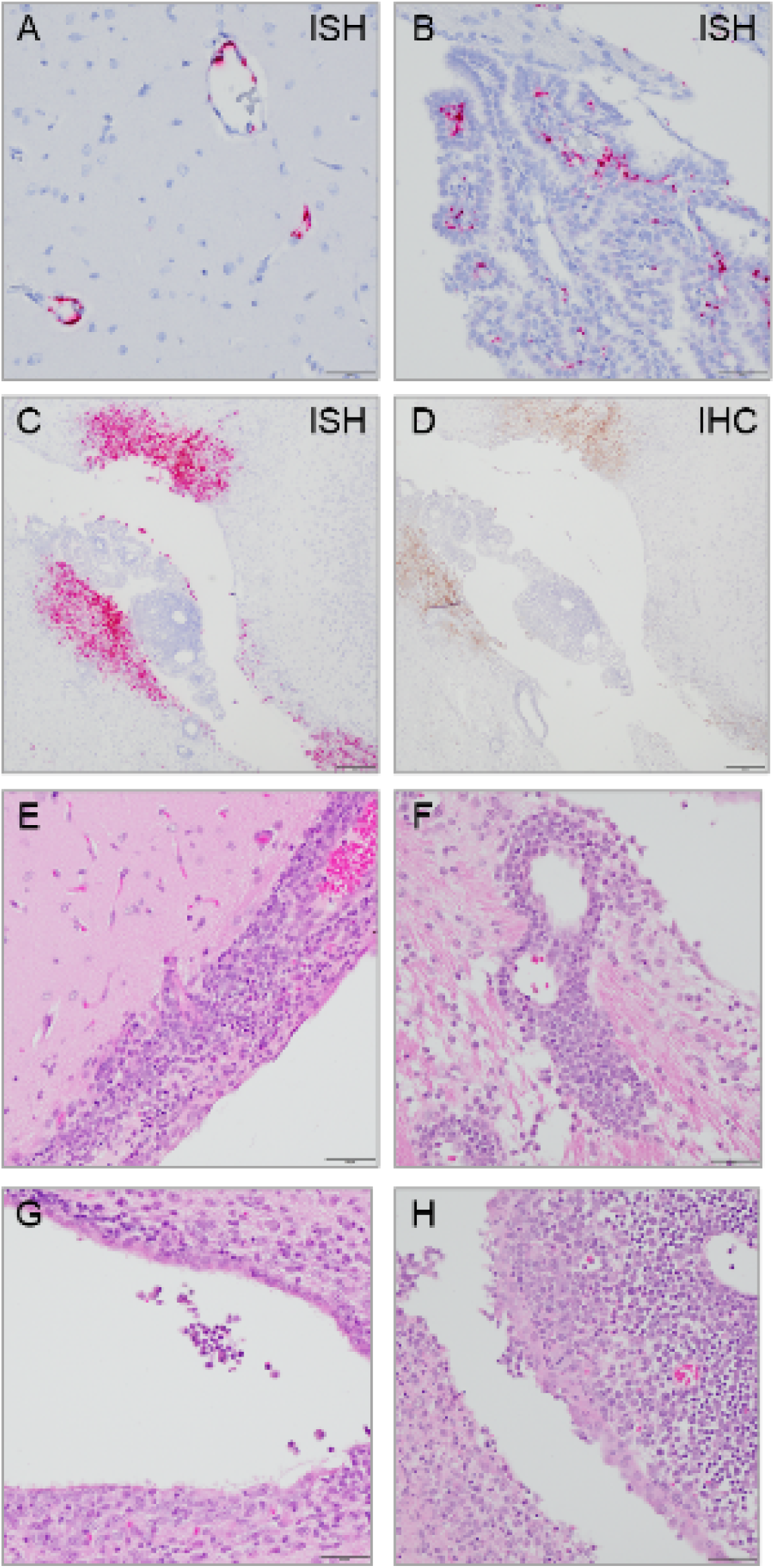
Brain histopathology in animals that died of atypical EVD. Ferret brain tissue sections stained by RNA in situ hybridization (ISH, red; A–C) or immunohistochemistry (IHC, brown; D) from an acute ferret (A, B) and an atypical ferret (C, D). EBOV genomic RNA (red) was only detected in endothelial vessels in the brain (A, B) of the acute ferret. Both EBOV genomic RNA (red; C) and antigen (brown; D) were detected in the infiltrates of meninges and ventricular system of the atypical animal. Nuclei were counterstained blue with hematoxylin. Representative brain tissue sections stained with hematoxylin and eosin (H&E) from an atypical ferret, revealing meningoencephalitis (E, F), ventriculitis (G), and choroid plexitis (H).

Inflammation of the brain was further substantiated by transcriptomic and proteomic analyses, which revealed variable expression and abundance of genes and proteins associated with a range of biological processes in a subset of ferrets from the second challenge experiment (Fig. 4). Of the fifty named genes with the most variable expression, many were associated with immune system processes and inflammatory responses, defence responses, and programmed cell death, as well as various cellular signalling, differentiation, and transport processes; however, none was differentially expressed only in animals that succumbed (Fig. 4A). Among the fifty most variably expressed genes overall, a cluster of eleven unnamed genes with no known functions was found to be significantly upregulated only in animals that experienced atypical disease (Fig. S3). While one of these genes had no known homologues, the other ten genes had homologues in other species (50-90% similarity) and were all identified as immunoglobulin-like domain-containing proteins.

**Figure 4.**
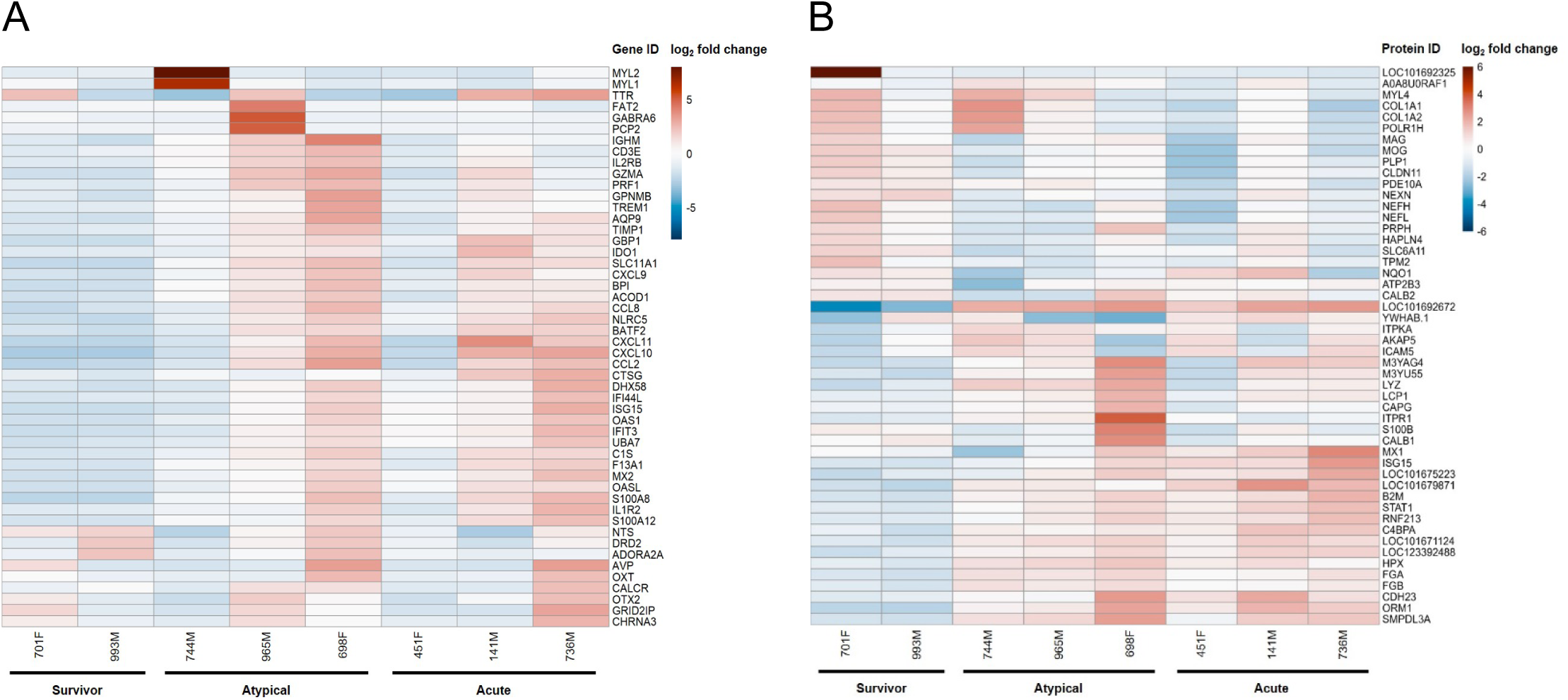
Transcriptomic and proteomic analyses of ferret brain tissue. Brain tissue samples from a subset of ferrets (all from the second challenge experiment) were subjected to transcriptomic and proteomic analyses. A heatmap from the transcriptomic analysis shows the differential expression of the fifty named genes with the most variable expression, with log_2_ fold change centered for each gene (A). A heatmap from the proteomic analysis shows the differential abundance of the fifty proteins with the highest variance among all animals, with log_2_ fold change centered for each protein (B).

Proteomic analysis complemented the transcriptomic findings and identified many of the same biological processes (Fig. 4B). The fifty most variably abundant proteins fell into two main clusters: those higher in survivors compared to non-survivors and those higher in non-survivors compared to survivors. Proteins more abundant in survivors were associated with processes related to recovery from acute illness, such as anatomical structure development, signaling, cell differentiation, adhesion and motility, and cellular transport. Conversely, proteins found more abundantly in acute or atypical animals were associated with processes involved in the immune and nervous systems, defence responses, and programmed cell death, as well as cellular signaling, differentiation, and transport processes. Notably, haptoglobin (LOC101692672), which binds free hemoglobin resulting from the lysis of red blood cells, was found in much higher abundance in the brains of acute and atypical animals than survivors, suggesting intracerebral hemorrhage and/or a break down in the blood-brain barrier (29). Moreover, several proteins associated with nervous system processes, namely ITPR1, S100B, CALB1, and CDH23, were more abundant in the brains of atypical animals, especially 698F, in concordance with the apparent neurological nature of disease that these animals experienced.

### Blood chemistry in ferrets that died of atypical disease was mostly normal

Dramatic changes in certain blood chemistry parameters are hallmarks of EVD and generally indicate organ damage caused by infection. In the ferrets that died of atypical EVD, however, the majority of these parameters were normal (Fig. 5 and Fig. S4). Alanine aminotransferase (ALT), a marker of liver function, was increased in all acute animals, as well as several atypical and surviving animals, during the week following inoculation (Fig. 5A). These levels decreased in atypical and surviving animals around 9 DPI and remained within their normal range for the rest of the study. Thus, at the terminal time points, the mean level of ALT present in the atypical animals was significantly lower than that in the acute animals and indistinguishable from the ALT level in the survivors (Fig. 5B). Two other markers of liver function—alkaline phosphatase (ALP) and total bilirubin (TBIL)—showed a similar trend. ALP and TBIL levels were elevated in most acute animals shortly after inoculation but remained low in both the atypical and surviving animals (Fig. 5C, E). At the terminal time points, mean levels of ALP and TBIL were significantly lower in the atypical animals compared to the acute animals, once again resembling what was observed in the surviving animals at the end of the study (Fig. 5D, F). The same trends were seen for amylase, blood urea nitrogen (BUN), and K^+^ (Fig. S4A-F). Albumin levels decreased immediately following inoculation in most animals, but most dramatically in those that died of acute disease (Fig. 5G). Levels rebounded in both the atypical and surviving animals such that, at terminal time points, the mean level of albumin in the atypical animals was significantly higher than the mean level in the acute animals (Fig. 5H). Likewise, Ca^2+^ levels in these animals followed the same general pattern (Fig. S4G, H).

**Figure 5.**
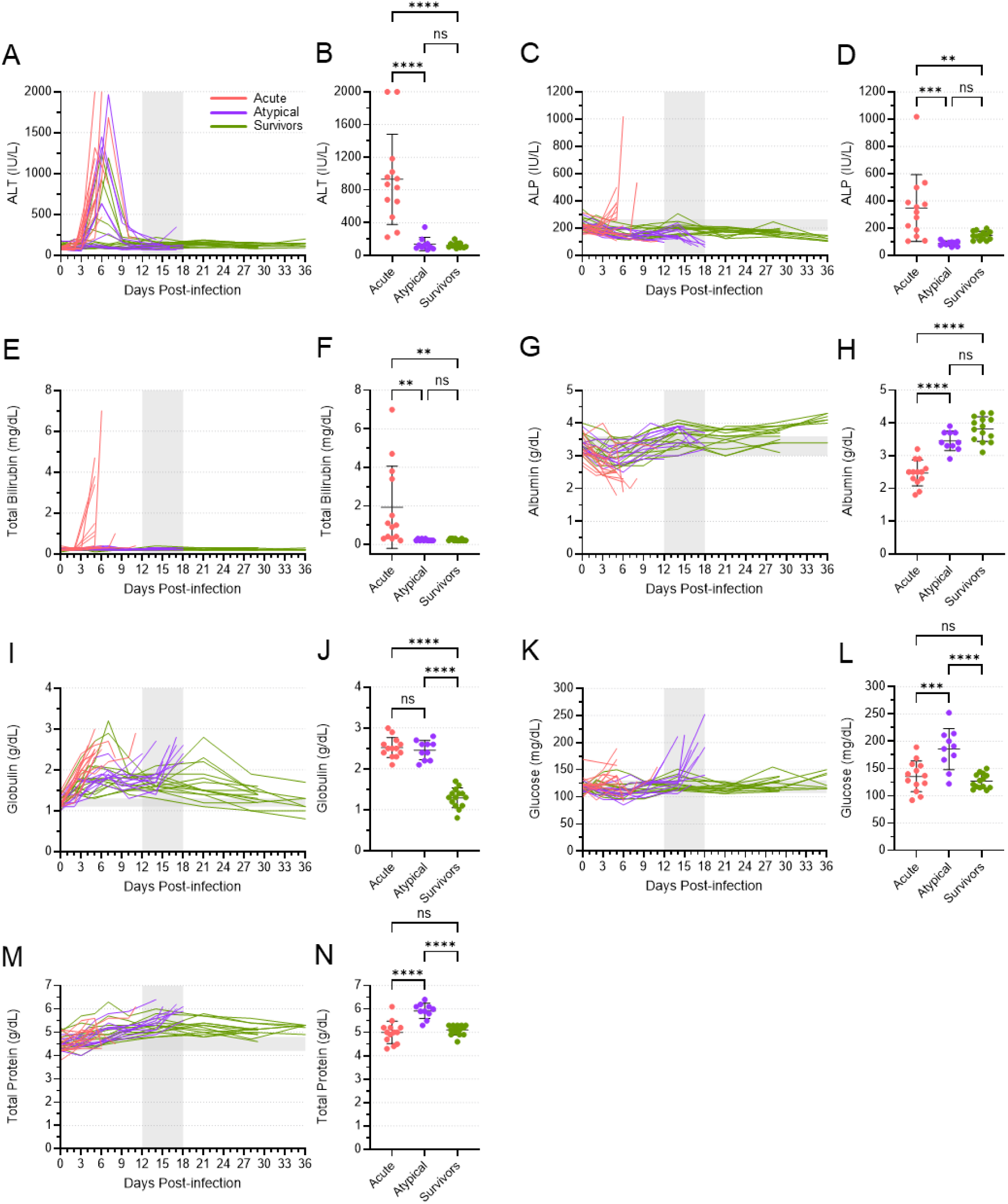
Blood biochemistry parameters for acute, atypical, and surviving ferrets. Levels of alanine aminotransferase (ALT) (A, B), alkaline phosphatase (ALP) (C,D), total bilirubin (E, F), albumin (G, H), globulin (I, J), glucose (K, L), and total protein (M, N) were quantified in all blood samples from all animals. Levels of each of these analytes are depicted over time (A, C, E, G, I, K, M). The vertical area shaded grey, from 12 to 18 DPI, represents the window in which atypical disease was observed, while the horizontal area shaded grey represents the normal range for each parameter. The levels of each analyte at the terminal time points are also depicted separately (B, D, F, H, J, L, N), with means and standard deviations indicated. Mean levels were compared using a one-way ANOVA with Tukey’s multiple comparison test. ns, not significant; **, p≤0.01; ***, p≤0.001; ****, p≤0.0001.

Not all blood chemistry parameters in the atypical animals resembled the survivors. While globulin levels increased in many animals immediately following inoculation, they remained high in the acute animals, which succumbed around the same time that globulin levels peaked (Fig. 5I). Levels subsequently decreased in atypical and surviving animals, only to increase again in the atypical animals at the time of death. Thus, at the terminal time points, the levels of globulin in the acute and atypical animals were equivalent and significantly higher than what was observed in the surviving animals (Fig. 5J). A similar trend, albeit less well pronounced, was observed for Na^+^ levels (Fig S4I, J). Conversely, the changes in glucose and total protein levels in the atypical animals differed from both the acute and surviving animals. Glucose levels increased significantly at the time that animals succumbed to atypical disease, while they remained relatively normal in both acute and surviving animals at their terminal time points (Fig. 5K, L). Total protein levels increased gradually for all animals after inoculation, peaking with the atypical animals at their time of death and then decreasing again in the surviving animals (Fig. 5M, N). These data further suggest that the atypical disease observed in some ferrets was distinct from acute EVD.

### Hematology parameters differed in animals that died of atypical disease

Changes in the cellular composition of the blood of the atypical ferrets also differed from the changes observed in both the acute animals and survivors (Fig. 6). A drop in lymphocyte numbers was observed in all animals around 3 DPI, which is typical for EVD (Fig. 6A). While most of the acute animals died shortly thereafter with low numbers of lymphocytes in their blood, numbers rebounded in both the atypical and surviving animals, with some of these animals showing dramatic increases in their lymphocyte counts around 7 DPI. Lymphocyte numbers eventually returned to normal in the surviving animals, but they dropped once again in the atypical animals at the time of death. Thus, at the terminal time points, both the acute and atypical animals had similar mean lymphocyte counts that were significantly lower than what was observed in the surviving animals (Fig. 6B). Conversely, neutrophil numbers trended upwards for all animals following infection, with several atypical animals showing high peak numbers between 6 and 10 DPI (Fig. 6C). Numbers decreased in most surviving and atypical animals before increasing again in many atypical animals or eventually returning to normal levels in the surviving animals. As a result, at the terminal time points, atypical animals showed mean neutrophil numbers that were higher than both acute and surviving animals, although the difference between that atypical and acute levels was not statistically significant (Fig. 6D). Similarly, the mean number of monocytes in the atypical animals at the terminal time points was significantly higher than that of both the acute and surviving animals (Fig. 6E, F). Prior to the terminal time points, there was an early peak in monocyte numbers in most animals around days 2 and 3, followed by a second, more dramatic peak in some of the atypical animals around days 7 to 9 (Fig. 6E). Monocyte numbers remained relatively high or increased around the time that the atypical animals succumbed (Fig. 6F). Total white blood cell counts reflected the sum of the changes in lymphocyte, neutrophil, and monocyte counts (Fig. 6G). A general decrease in numbers was evident within the first 3 to 6 days after inoculation, perhaps driven by concomitant drop in lymphocytes. A peak around 7 DPI in many atypical and surviving animals aligned with similar peaks in lymphocyte and neutrophil numbers; this peak was followed by a return to normal or near normal white blood cell numbers for most remaining animals. Accordingly, no differences in the mean numbers of white blood cells were observed among acute, atypical, and surviving animals at their terminal time points (Fig. 6H). Platelet counts varied considerably throughout the study, although the numbers in the acute animals trended downwards at the terminal time points (Fig. 6I). Platelet numbers in the atypical animals also trended downwards at or near the time of death, but the mean numbers were still significantly higher than what was observed in the acute animals (Fig. 6J). The surviving animals also showed variability in their platelet counts, but they returned to the normal range at study endpoint, when the mean level was significantly higher than both the acute and atypical animals at their endpoints (Fig. 6J).

**Figure 6.**
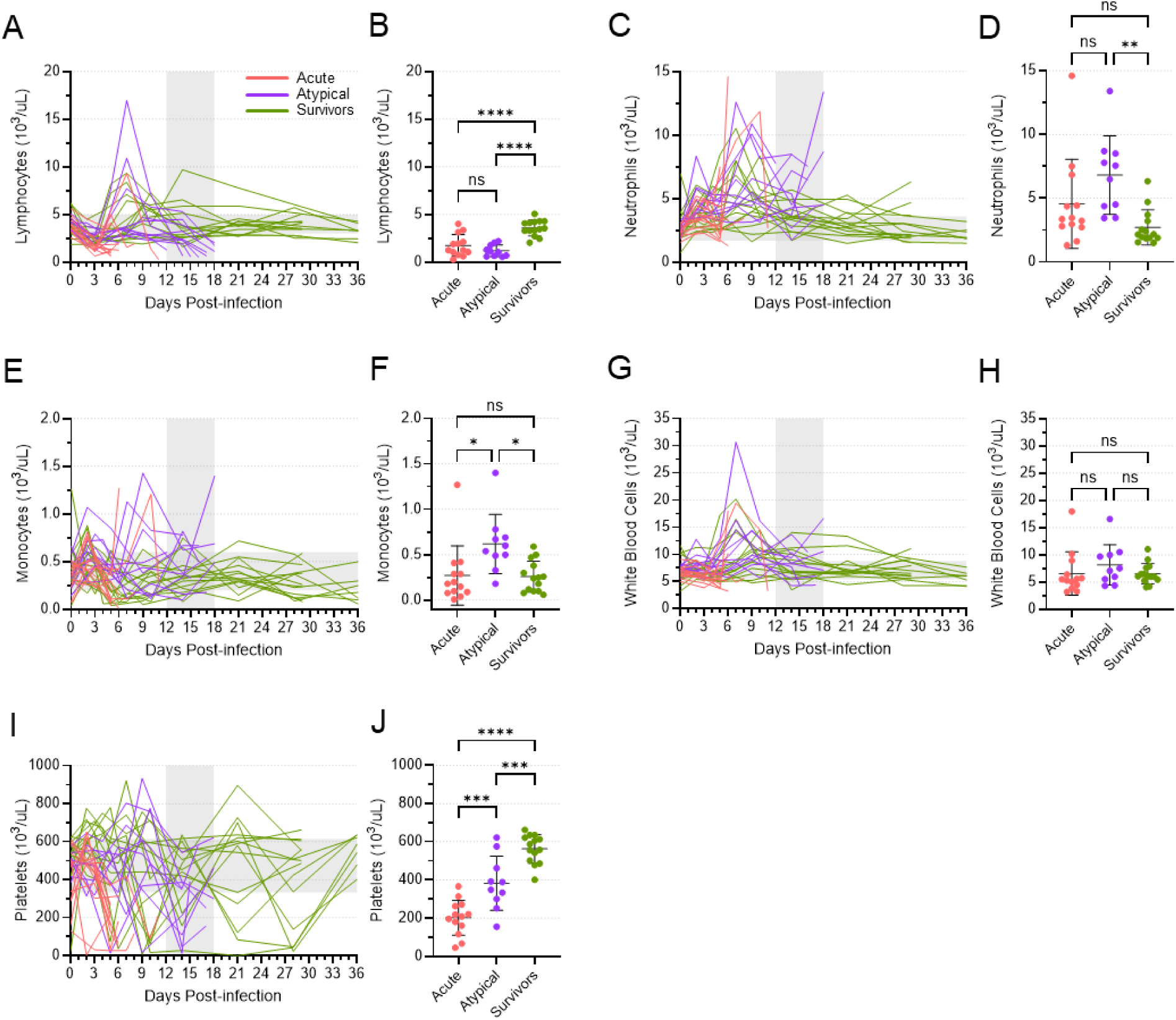
Blood cell counts for acute, atypical, and surviving ferrets. Lymphocytes (A, B), neutrophils (C, D), monocytes (E, F), white blood cells (G, H), and platelets (I, J) were enumerated in all blood samples from all animals. Cell counts are depicted over time (A, C, E, G, I). The vertical area shaded grey, from 12 to 18 DPI, represents the window in which atypical disease was observed, while the horizontal area shaded grey represents the normal range for each cell type. Cell counts at the terminal time points are also depicted separately (B, D, F, H, J), with means and standard deviations indicated. Mean levels were compared using a one-way ANOVA with Tukey’s multiple comparison test. ns, not significant; *, p≤0.05; **, p≤0.01; ***, p≤0.001; ****, p≤0.0001.

### Differential cytokine responses were induced during acute and atypical disease

The levels of twelve different cytokines increased in nearly all animals over the first 9-10 days following virus inoculation, with the most dramatic increases observed in the animals that died of acute EVD (Fig. 7 and Fig. S5). In almost all cases, cytokine levels peaked in acute animals at their terminal time point, between 5 and 8 DPI, with IP-10, MCP-1, MIP-1β, IL-6, IL-12p40, and IL-8 reaching particularly high levels (Fig. 7). The cytokine expression patterns in the atypical and surviving animals were generally similar to each other, with peak levels reached around 7-9 DPI, after which they typically declined. Interestingly, however, increases in some cytokines—namely IP-10, MCP-1, MIP-1β, IL-6, and to a lesser extent, IL-4—coincided with the presentation of atypical EVD (Fig. 7). With the exceptions of MIP-1β and IL-4, the mean levels of these cytokines in the atypical animals at their terminal time points were significantly higher than what was observed in the surviving animals, although they were also significantly less than what was observed in the acute animals. These data once again demonstrate a stark difference in the disease phenotype between animals that succumbed to acute EVD and those that succumbed to atypical EVD.

**Figure 7.**
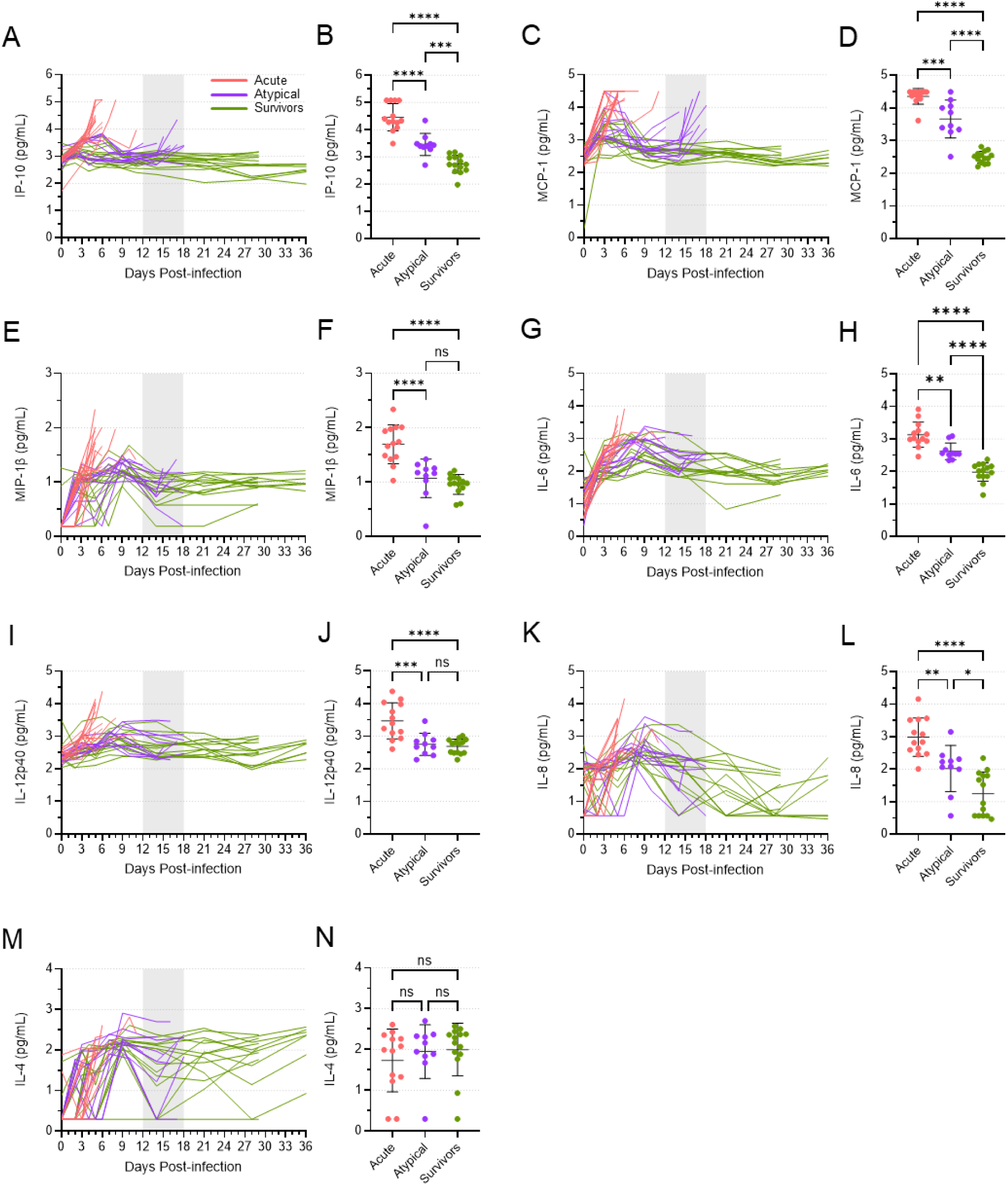
The cytokine response in acute, atypical, and surviving ferrets. Levels of IP-10 (A, B), MCP-1 (C, D), MIP-1β (E, F), IL-6 (G, H), IL-12p40 (I, J), IL-8 (K, L), and IL-4 (M, N) were quantified in all serum samples from all animals via Luminex assay. Levels of each cytokine are depicted over time (A, C, E, G, I, K, M). The vertical area shaded grey, from 12 to 18 DPI, represents the window in which atypical disease was observed. The levels of each cytokine at the terminal time points are also depicted separately (B, D, F, H J, L, N), with means and standard deviations indicated. Mean levels were compared using a one-way ANOVA with Tukey’s multiple comparison test. ns, not significant; *, p≤0.05; **, p≤0.01; ***, p≤0.001; ****, p≤0.0001.

### Both atypical and surviving animals mounted a humoral immune response

To understand the humoral immune response mounted by infected animals, we quantified the amount of EBOV GP-specific IgG present in the plasma of atypical and surviving animals, beginning with samples collected on 9 DPI (Fig. 8A). All animals exhibited a detectable IgG response against EBOV GP, with endpoint titers ranging from 1:800 to as high as 1:25,600. In general, IgG levels increased over time, as expected based on the continued maturation of the humoral immune response. Although the geometric mean IgG endpoint titer for the survivors was significantly higher than that of the atypical animals at the terminal time points (Fig. 8B), this can likely be attributed to the difference in time between when the terminal samples were collected (12-18 DPI for the atypical animals and 29 or 36 DPI for the surviving animals). Indeed, comparing the mean titer of the atypical animals at their terminal timepoints with the mean titer of the survivors at 14 DPI revealed no significant difference (Fig. 8B). Thus, both atypical and surviving animals appeared to mount a humoral immune response of equivalent magnitude, making it unlikely that this response was a factor in the development of atypical EVD.

**Figure 8.**
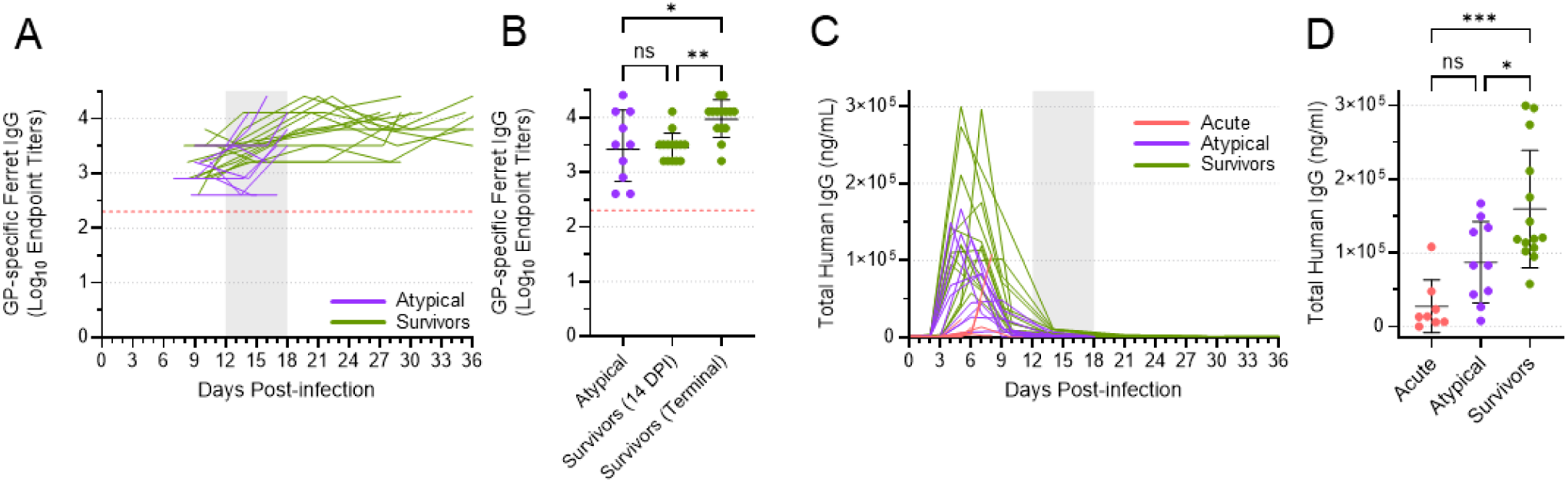
Antibody quantification in acute, atypical, and surviving ferrets. Levels of EBOV GP-specific IgG were determined via ELISA (A, B) in all serum samples from the atypical and surviving animals, starting with samples collected on 9 DPI. Levels of antibody are depicted over time (A), with each line in the graph off-set slightly relative to the x-axis to ensure that data from individual animals could be discriminated. EBOV GP-specific IgG levels at the terminal time points, as well as at 14 DPI for the surviving animals, are also depicted separately (B), with means and standard deviations indicated. The red dotted line indicates the lower limit of quantification for this assay. Total levels of human IgG were determined via ELISA (C, D) in all serum samples from all animals. Levels of antibody are depicted over time (C). Human IgG levels at the terminal time points are also depicted separately (D), with means and standard deviations indicated. The vertical area shaded grey, from 12 DPI to 18 DPI, in (A) and (C) represents the window in which atypical disease was observed. Mean levels in (B) and (C) were compared using a one-way ANOVA with Tukey’s multiple comparison test. ns, not significant; *, p≤0.05; **, p≤0.01; ***, p≤0.001.

### Levels of circulating human mAbs differed among acute, atypical, and surviving animals

To better understand the dynamics of human mAb concentration following administration, we used an ELISA specific for human IgG to quantify the total amount of human antibody present in all ferret plasma samples (Fig. 8C). Human antibody levels peaked between 3 and 7 DPI, following administration of the antibodies between 2 and 6 DPI. By 14 DPI, human antibody levels were universally very low, and they effectively disappeared by the terminal time points in the atypical and surviving animals.

Comparing the peak levels of human antibody in the acute, atypical, and surviving animals revealed significant differences (Fig. 8D). The animals that died of acute EVD exhibited very low to undetectable levels of human antibody in most samples, with a mean peak level that was significantly less than what was observed in the survivors. The mean peak level of human antibody in the animals that died of atypical disease was higher than that observed in the acute animals, which died earlier, but significantly lower than the mean peak level observed in the animals that survived. These data suggest that there were differing total levels of human mAb circulating in each ferret, and that the peak levels may correlate with disease outcome. However, caution is required in interpreting these data since the concentration of human antibody detected in each serum sample is likely affected by a number of different factors, including antibody uptake following intraperitoneal injection and antibody half-life, the latter of which may be further influenced by the binding of these antibodies to the virus and host responses against the foreign human antibodies.

To determine the half-lives of the human mAbs in ferrets, we conducted a pharmacokinetic experiment in a group of uninfected animals. Ferrets were injected with 30 mg/kg of one of two human IgG1 antibodies, either 1C3 or 1C11, and serum samples were collected regularly up to 504 hours post-treatment. Levels of each antibody were then quantified using the same human IgG ELISA as described above, and the half-life of each antibody in each animal was calculated (Fig. S6). The half-lives of these antibodies proved to be strikingly short in the ferrets, with 1C3 ranging from 29.76 hours to 78.41 hours (mean 50.2 hours) and 1C11 ranging from 33.51 hours to 65.64 hours (mean 54.85 hours). It seems likely that such short half-lives might have contributed to the reduced efficacy of these mAbs in the ferret model.

## DISCUSSION

The shortage of NHPs for research—not to mention the high costs and extensive ethical considerations—has inspired examination of alternate animals that might serve as models for human disease and countermeasure evaluation. In this study, we set out to evaluate a human antibody cocktail with demonstrated efficacy in NHP models of EBOV and the related Sudan virus (SUDV) (30) to determine how a ferret model would serve in comparison. Surprisingly, most of the ferrets that died did not succumb to disease that resembled acute EVD. Rather, these animals remained healthy until relatively late post-infection, at which point their condition deteriorated rapidly and clinical signs consistent with neurological problems became evident. The pattern of dysregulated blood biochemistry typical of acute EVD was not present in these animals at the time of death, suggesting that most of the animals’ organ systems were functioning normally. Although a moderate pro-inflammatory cytokine response was observed in the blood of these animals just prior to death—coincident with an overall increase in neutrophils and monocytes and a decrease in lymphocytes—this immune response was significantly lower in magnitude compared to what was observed during acute EVD. While the majority of the atypical animals had no detectable viral RNA in their blood at 14 DPI, low levels of viral RNA reappeared at the terminal time points. Low levels of viral RNA were also noted in the livers and spleens of these animals, in contrast to the brain, which harboured very high levels of viral RNA in addition to infectious virus. Evidence of this RNA, as well as virus antigen, was also detected in the brain, via ISH and IHC, alongside histological evidence of meningoencephalitis, ventriculitis, and choroid plexitis. The host response in the brain, as assessed by transcriptomics and proteomics, further suggested significant pathological changes in the central nervous system, including a breakdown in the blood-brain barrier. Together, the unusual constellation of disease parameters that we observed in these animals formed a pattern distinct from that seen in animals that succumbed to acute disease, as well as those that survived.

To our knowledge, this is the first time that organ-specific persistence followed by fatal EVD recrudescence has been reported in ferrets; however, there are a handful of reports describing similar phenomena in NHPs (17). We previously described the case of a single rhesus macaque inoculated with EBOV and then treated with two doses of a mAb cocktail consisting of FVM04 and CA45 (25). This animal recovered from an apparently mild course of acute EVD only to develop severe atypical EVD approximately two weeks later, beginning on 21 DPI, and ultimately meeting the conditions for euthanasia on 26 DPI, with signs of disease consistent with meningoencephalitis. This animal had no detectable viral RNA in the blood at the time of death despite abundant levels in most tissues, including the brain, which was one of the only tissues from which infectious virus could be isolated. A subsequent case report described a similar story in which a rhesus macaque that had been vaccinated with the rVSV-EBOV vaccine and challenged with EBOV developed severe neurological signs of disease and respiratory distress at 21 DPI before requiring euthanasia on 28 DPI (28). As in the previous case, the manifestation of atypical EVD in this animal was not accompanied by viremia, although virus was detected in numerous immune-privileged tissues, including the brain. An earlier study described another instance of a cynomolgus macaque—this time treated with adenovirus-vectored IFN followed by two doses of the ZMAb antibody cocktail—that recovered from acute disease caused by EBOV before developing signs of disease around 22 DPI that were consistent with neurological involvement (26, 31). Although viremia had disappeared from this animal by 16 DPI, it rebounded in parallel with the manifestation of atypical EVD, with high levels of viral RNA found in both the blood and cerebral spinal fluid at the time of euthanasia on 25 DPI. Additionally, a report by Larsen et al. from 2007 describes atypical EVD in six rhesus macaques that had been challenged with EBOV and treated with various different therapeutics, including recombinant nematode anticoagulant protein c2, recombinant human activated protein C, recombinant human interferon-1a, and in one case, mAb KZ52 (27). Of the six animals, four survived longer than 20 DPI and appeared to recover from acute EVD before their clinical condition deteriorated rapidly, eliciting death or euthanasia. Notably, signs of neurological disorder were observed in three of these four animals, accompanied by relatively high levels of infectious virus in the brain and lower or absent levels of virus in common target organs, like the liver and spleen. Although the details differ slightly among each of these case reports, they all evoke a similar pattern: a rapidly progressing, late-onset disease that often includes neurological signs, along with abundant levels of virus in the brain, but often lacks most hallmarks of acute EVD, including prominent viremia, as well as marked disturbances in blood chemistry, hematology, and the innate immune response. This same pattern applies to the ferrets described in the present study, suggesting that ferrets, like NHPs, may also recapitulate less common aspects of EVD, at least under specific experimental conditions.

Three human cases of EVD recrudescence leading to meningoencephalitis have also been well-described in the literature, with symptoms reappearing 4-11 months following recovery from acute disease and culminating in the death of two of the three patients (15, 23). In all three cases, viral RNA was detected at much higher levels in the cerebral spinal fluid than in the blood, reflecting observations in ferrets from the present study as well as several of the NHP case studies. In a fourth case of EVD recrudescence in a human, which arose ∼5 months after initial recovery, meningoencephalitis was not described, with the patient instead presenting with signs of disease similar to what is observed in acute EVD (24). During their initial disease course, all four patients had been treated with a mAb therapeutic in conjunction with supportive treatment, while two of the patients also received post-exposure vaccination with rVSV-EBOV (23), and another one received the small molecule drug brincidofivir and convalescent plasma (15). The advanced level of care received by these four patients undoubtedly contributed to their survival from acute EVD.

While the mechanisms that underlie the development of atypical EVD—whether in NHPs, ferrets, or humans—are still being investigated, two retrospective histopathological studies on banked NHP samples have provided vital insight (32, 33). In 2017, Zeng et al. demonstrated persistent EBOV infection in the brains, eyes, and testes of a minority of rhesus macaques that had survived EBOV challenge (33). EBOV was found to persist within CD68+ monocytes/macrophages, and it was suggested that infection could disseminate progressively from the vascular structures associated with these tissues into the tissues themselves, causing an inflammatory immune response and potentially setting the stage for disease relapse. In 2022, the same group demonstrated that EBOV preferentially persists in the brain ventricular system of rhesus macaques that had been treated with a mAb therapeutic (32). CD68+ monocytes/macrophages were again identified as the cellular reservoir of persistent EBOV infection, and EBOV persistence was accompanied by ventriculitis, choroid plexitis, and meningoencephalitis. Of the 50 animals assessed, 8 showed evidence of EBOV persistence in the brain, and two of those animals went on to develop fatal recrudescence that followed the same pattern of atypical EVD, with EBOV infection and associated pathology essentially confined to the brain in both cases.

Although mAb treatment has been associated with instances of EVD relapse or recrudescence in animal models and humans (15, 23–25, 27, 28, 31), we recognize the circumstantial nature of this evidence. In particular, we are cognizant of survivorship bias, which might erroneously lead to the interpretation that mAb treatment causes recrudescence, rather than the interpretation that mAb treatment causes survival, which then creates the opportunity to observe recrudescence. Indeed, one cannot model recrudescence in an animal that does not survive acute infection. Since mAb therapeutics are among the most reliably successful treatment strategies for EVD (34), they may simply provide a means by which EBOV-infected animals can survive. Although mAb treatment often leads to virus clearance in these survivors, in some instances—perhaps when only low levels of circulating mAb are achieved—virus replication may be reduced but not eliminated, permitting EBOV to establish a persistent infection without causing acute disease and thereby creating the conditions for potential organ-specific recrudescence in immune-privileged sites. We also emphasize the fact that atypical EVD cannot be causally linked to a specific mAb therapeutic or to mAb therapeutics in general, given that other instances of recrudescence in humans and animals have also been associated with the administration of additional treatments, including post-exposure vaccination and small molecule drugs. The fact that atypical EVD has been observed in a vaccinated NHP (28), as well as NHPs that had been treated with non-mAb countermeasures (27, 33), supports the notion that atypical EVD may result from a variety of complex factors.

With the preceding caveats in mind, we envision a scenario that may explain the atypical disease we observed in ferrets in this study. EBOV is exceptionally virulent in these animals, resulting in high levels of virus replication in many tissues, including immune-privileged sites like the eyes and testes, as observed by Watson et al. (35), and the brain, as observed in this study. mAb treatment is highly effective at neutralizing virus, particularly in the circulation, where the highest concentration of mAbs is likely to exist, and in tissues that are readily permeated by antibodies, such as the liver and spleen (36). However, in immune-privileged tissues, mAb penetration is likely much lower, and, in the brain, it has been estimated to be just ∼0.35% of the plasma antibody concentration (36). Thus, while mAb treatment was likely able to eliminate virus from circulation, as well as several EBOV target organs, and prevent acute EVD, it may have been unable to eliminate EBOV from the brain, where virus persisted for approximately two weeks before re-emerging primarily as a brain-specific inflammatory disease. This is further supported by transcriptomic and proteomic markers of inflammation that were identified in the brains of atypical animals (as well as those of acute animals), with additional evidence of damage to the blood-brain and blood-CSF barriers seen in the enrichment of genes and proteins—such as haptoglobin and immunoglobulin-like domain containing proteins—that would normally be found only in the periphery. In line with these data, atypical EVD in the ferrets was also characterized by meningoencephalitis, ventriculitis, and choroid plexitis, reflecting the pathology observed in NHPs that experienced atypical EVD (32, 33). Whether EBOV also persisted in ferret CD68+ monocytes/macrophages, as was observed in the NHPs, remains to be determined. It is important to note, however, that while we have convincing evidence pointing to infection in the brains of these animals, as well as a lack of infection in the liver and spleen, we cannot rule out the possibility that EBOV persisted elsewhere in these ferrets and re-emerged from these sites after the therapeutic antibodies disappeared from circulation. Although EBOV infection and inflammation in the brain is consistent with the neurological signs of disease that were also observed at the terminal time points, we appreciate the fact that, in ferrets, neurological disease can be difficult to distinguish from generalized weakness (37). This makes it difficult to state conclusively that the manifestation of atypical EVD was, in fact, specifically neurological and not systemic.

An additional factor that likely had a significant impact on the development of atypical EVD in these ferrets is antibody pharmacokinetics. The half-life of human mAbs in ferrets is known to be remarkably short, with previous studies giving half lives of ∼30 to ∼70 hours (38). Our analysis of the two treatment mAbs used in this study matches the previous values, with estimated half lives of ∼50 to 55 hours. These short half lives contrast dramatically with what has been observed for mAbs in humans and NHPs. For instance, the three mAbs in the REGN-EB3 cocktail, which has been approved for the treatment of EVD, have reported half lives that range from 8-16 days in cynomolgus macaques and 10-14 days in rhesus macaques (39), while in humans, the half lives range from ∼22-27 days (40). Similarly, mAb114, another approved mAb therapeutic for treating EVD, has a half life of 7-15 days in rhesus macaques and over 24 days in humans (41). Thus, the extremely short half-lives of human mAbs in ferrets likely reduces treatment efficacy as compared to humans and NHPs. In practice, this means that the concentrations of human antibodies in EBOV-infected ferrets would be expected to decrease rapidly post-treatment, as the antibodies are eliminated by the ferret or consumed via binding to virus. Indeed, we observed that the two treatment mAbs essentially disappeared from ferret circulation between 10 and 14 DPI, after which atypical EVD began to manifest in some animals. Moreover, the animals that would eventually go on to develop atypical EVD exhibited significantly lower peak levels of mAb in their blood than the survivors. Thus, in animals that experienced atypical EVD, lower levels of mAb may have been less effective at completely eliminating virus, which then rebounded to cause disease after the antibodies disappeared from circulation.

Regardless of the mechanisms underlying the development of atypical EVD, our study suggests that ferrets may be a suitable model for better understanding this rare consequence of EBOV infection, particularly since ferret experiments can be performed with larger numbers of animals than NHP experiments. Our in-depth characterization of this disease revealed a distinct pattern of virus replication and host response biomarkers that reflects similar observations in NHPs, which are considered the gold-standard animal model for filovirus infections. Future work will focus on further refining and characterizing this model system, with the intention of answering several key questions. For example, whether infection with other orthoebolaviruses results in atypical disease is unclear; however, a recent report demonstrating SUDV persistence in immune-privileged organs of NHPs suggests that this may be possibility in ferrets as well (42). It would also be interesting to know whether the incidence of atypical EVD following mAb treatment can be reduced, perhaps by delivering an additional mAb dose later post-infection or by co-administering a small molecule therapeutic alongside mAb treatment, which has recently proven successful in treating SUDV and the related filovirus, Marburg virus (43, 44).

Whether ferrets are a useful model, in general, for evaluating the efficacy of EBOV countermeasures, such as mAbs, also remains an open question. On the one hand, ferrets are susceptible to disease caused by wild type EBOV, and they exhibit many of the clinical features of EVD observed in humans and NHPs (21). On the other hand, EVD in ferrets is severe and rapid, setting a potentially very high bar for therapeutic success. With respect to mAb evaluation, our data suggest that the most successful treatment regimen involves inoculating animals via the IN route and treating them early. In addition to representing a more natural route of infection, mucosal inoculation may result in slower virus replication kinetics compared to IM inoculation, therefore increasing the effective treatment window. Delivering mAbs early within this window may then reduce virus replication quickly, before it becomes uncontrollable. Another factor to consider when evaluating mAbs in ferrets is their pharmacokinetics. The short half-lives of human mAbs in ferrets, especially compared to humans and NHPs, would be expected to significantly impact their efficacy, and differences in ferret Fc receptors may mitigate antibody effector functions. Thus, evaluation of mAbs in ferrets may only be useful for antibodies that primarily function by mechanical neutralization, as opposed to Fc-receptor mediated protection. Alternatively, one may need to explore chimerizing the antibodies to present a natural ferret Fc.

EBOV persistence carries with it significant public health implications. The sporadic re-emergence of virus from immune privileged compartments not only threatens the lives of otherwise recovered EVD patients, but it also threatens the lives of others in the community, re-igniting waning outbreaks or inciting new ones, sometimes years later (6, 8, 14, 23, 45, 46). Unfortunately, we lack a robust and practical model system in which we can consistently recapitulate virus persistence and recrudescence. While they are not perfect model systems, ferrets may have a significant role to play in the continued investigation and characterization of these unique phenomena. Our work here lays the groundwork for using EBOV-infected ferrets in conjunction with post-exposure treatment to reproducibly generate atypical EVD outcomes that closely resembles the brain-specific recrudescent disease observed in both NHPs and humans.

## MATERIALS AND METHODS

### Animal Ethics and Biosafety Statement

All animal work involving infectious filoviruses was conducted in the containment level 4 facility at the Canadian Science Center for Human and Animal Health (CSCHAH) in Winnipeg, Canada. All animal experiment protocols were reviewed and approved by the Animal Care Committee at the CSCHAH, in accordance with guidelines from the Canadian Council on Animal Care (CCAC). All staff working on animal experiments completed education and training programs according to the standard protocols appropriate for this level of biosafety.

### Viruses and Cells

EBOV variant Makona C07 (H.sapiens-wt/GIN/2014/Makona-Gueckedou-C07; GenBank accession no. KJ660347.2) was used as the challenge virus in this study. The virus stock was mycoplasma negative, sequence confirmed, and stored at -80°C. Vero E6 cells were obtained from the American Type Culture Collection and maintained at 37°C and 5% CO_2_ in Dulbecco’s modified Eagle’s medium (DMEM; HyClone) supplemented with 5% heat-inactivated fetal bovine serum (HyClone), 2 mM L-glutamine (HyClone), and 50 U/ml penicillin and 50 ug/ml streptomycin (HyClone).

### Ferret Challenge Studies

Two independent ferret challenge experiments were performed (Fig. S1) with domestic ferrets (*Mustela putorius furo*) purchased from Marshall BioResources (New York, USA), where they had been implanted with a temperature and ID transponder subcutaneously over the dorsal aspect of the caudal region. Upon arrival at CSCHAH, animals were acclimated for at least 7 days prior to the commencement of the experiment. Throughout the study period, animals were given food and water *ad libitum*, provided with environmental enrichment, and monitored at least twice daily.

For the first challenge experiment, twenty ferrets (10 male and 10 female; ∼8 weeks old) were randomly divided into one of two groups and inoculated with a target dose of 1000 TCID_50_ EBOV via the intramuscular (IM) or intranasal (IN) route (Table S1). Both inoculations were delivered in a total volume of 500 ul, split evenly across two sites in the rear quadriceps (IM) or between each nostril (IN). Following virus inoculation, eight animals (four from each group; equal males and females) were treated via intraperitoneal (IP) injections with 30 mg/kg each of 1C3 and 1C11 diluted in PBS on 2 and 5 DPI; another eight animals (four from each group; equal males and females) were treated with the same dose of mAb on 3 and 6 DPI. Four animals (two from each group; equal males and females) were treated with PBS only. For the second challenge experiment, seventeen ferrets (7 male and 10 female; ∼9 weeks old) were randomly divided into one of two groups and once again inoculated with EBOV via the IM or IN route, as described above (Table S1). Following virus inoculation, eight animals (four from each group; equal males and females) were treated via IP injections with 30 mg/kg each of 1C3 and 1C11 on 2 and 4 DPI, as above; another eight animals (four from each group; 3 males and 5 females) were treated with the same dose of mAb on 3 and 5 DPI. One animal (female) that had been inoculated via the IN route was treated with PBS only. Antibody doses were calculated separately for male and female animals, based on the average weight of all animals of each sex. Thus, for the first experiment, all females received 13.2 mg of each mAb, while all males received 15.9 mg of each mAb. For the second experiment all females received 15.4 mg of each mAb and all males received 18.7 mg. Animals were monitored daily for clinical signs of disease including changes in body weight, temperature (via transponder), physical activity, and food/water intake. Animals that reached the clinical scoring criteria for humane endpoint were euthanized. On predetermined days (-2, 2 or 3, 5 or 6, 9, 14, 21, and 29 DPI for the first experiment and -4, 2 or 3, 4 or 5, 7, 10, 14, 21, 28, and 36 DPI for the second) or when an animal reached the humane endpoint, animals were anesthetized and given a physical examination, including weight and rectal temperature measurements, by qualified study personnel. Whole blood (EDTA) and plasma (lithium heparin) samples were also collected during these examinations. Terminal blood samples were obtained from animals that reached the humane endpoint, after which the animals were euthanized and tissues (liver, spleen, brain) were collected. Note that, from the first experiment, liver and spleen samples were harvested for RNA extraction from all treated animals (but not the control animals), while brain samples were harvested from all animals that were euthanized after 12 DPI. No tissue samples were saved from this study for the isolation of infectious virus. From the second experiment, liver, spleen, and brain samples were harvested from all animals for both RNA extraction and virus isolation.

### Antibody Pharmacokinetics Study in Ferrets

Eight ferrets (four male and four female; ∼9 weeks old) were purchased and acclimated as described above. The animals were randomly divided into one of two groups and treated via IP injection with 30 mg/kg of 1C3 or 30 mg/kg of 1C11. Antibody doses were calculated based on the individual weight of each animal; the males therefore received 18-20.4 mg of mAb, while the females received 14.7-15.9 mg of mAb. Serum samples were obtained from each animal one day prior to mAb treatment and at regular intervals after mAb treatment: 0 (i.e., immediately after treatment), 1, 6, 24, 48, 72, 96, 120, 144, 168, 336, and 504 hours post-treatment.

### Viral RNA Extraction and Quantification

Viral RNA was extracted from EDTA blood samples using the QIAamp Viral RNA Mini Kit (Qiagen) following manufacturer’s instructions. Tissue samples were stored in RNAlater (ThermoFisher Scientific), after which RNA was extracted using the RNeasy Plus Mini Kit (Qiagen) according to manufacturer’s instructions. EBOV RNA levels were quantified via reverse transcription quantitative PCR (RT-qPCR) using the LightCycler 480 RNA Hydrolysis Probes kit (Roche) on a QuantStudio 3 Real-Time PCR System (Applied Biosystems) using the following EBOV-specific primers and probes: forward primer, 5’-CAGCCAGCAATTTCTTCCAT-3’; reverse primer, 5’-TTTCGGTTGCTGTTTCTGTG-3’; probe 1, 5′-6-FAM-ATCATTGGC/ZEN/GTACTGGAGGAGCAG-3IABkFQ-3′; and probe 2, 5′-6-FAM-TCATTGGCG/ZEN/TACTGGAGGAGCAGG-3IABkFQ-3’ (Integrated DNA Technologies). The thermocycling conditions were as follows: 63°C for 3 min, 95°C for 30 s, followed by 45 cycles of 95°C for 15 s and 60°C for 30 s. The cycle threshold (Ct) values were converted to genome equivalents per milliliter (GEQ/mL) or gram (GEQ/g) using standard curves generated from a plasmid encoding the EBOV polymerase (L) gene.

### Quantification of Infectious Virus

Whole blood and tissues samples that were positive for EBOV RNA via RT-qPCR were assessed for the presence of infectious virus. Tissue samples were homogenized in DMEM using 5-mm stainless steel beads in a Bead Ruptor Elite Tissue Homogenizer (Omni International, Inc.) at a frequency of 4 m/s. Homogenates were clarified by centrifugation at 1000 x g for 10 minutes, after which they were tenfold serially diluted from 10^-1^ to 10^-8^ in DMEM containing 2% FBS. Whole blood samples were tenfold serially diluted from 10^-2^ to 10^-9^. A volume of 200 μl of each dilution was then added to a confluent monolayer of Vero E6 cells in 96-well plates, in triplicate. The plates were incubated at 37°C with 5% CO_2_ and assessed on 13 DPI for cytopathic effect. The median tissue culture infectious dose (TCID_50_) result was calculated for each sample using the Reed-Muench method (47).

### Genome Sequencing

EBOV whole genome sequencing was performed as described previously (48). Briefly, extracted viral RNA samples (8 μl) were treated with ezDNAse (Invitrogen) according to the manufacturer’s instructions. RNA extracts were combined with 50 ng/μl random hexamer primers and 0.2 mM dNTPs and heated to 65°C for 5 min, after which the primers were annealed by cooling on ice for 1 min. The Superscript III First-Strand Synthesis System (Invitrogen) was used to carry out cDNA synthesis from the hexamer-annealed RNA, per manufacturer’s instructions. cDNA was synthesized at 55°C for 90 min followed by inactivation of enzymes at 85°C for 5 min. Second strand synthesis was performed at 16°C for 16 hours in a total of 40 μl reaction, which included 10X Second Strand Synthesis Buffer, 10 U DNA Polymerase I, 0.35 U *E. coli* RNase H, 1.25 U *E. coli* DNA Ligase, and 0.375 mM dNTP mixture (all reagents from New England Biolabs). Resulting dsDNA was used as template for the TruSeq DNA Nano LP kit (Illumina) following the manufacturer’s protocol.

A total of 316 120-nt custom biotinylated lockdown DNA probes covering the entire EBOV genome in a double tiling approach were designed based on GenBank accession no. KJ660347.2 and purchased from Integrated DNA Technologies (Coralville, IA, USA). For EBOV-specific library enrichment, equal volumes of up to eight final libraries were pooled with the lock-down probes, per the xGen hybridization capture of DNA libraries protocol Version 2 (Integrated DNA Technologies). Biotinylated probes hybridized to EBOV libraries were captured with Dynabeads™ M270-streptavidin magnetic beads (Invitrogen) and washed extensively to remove unbound nucleic acids, following the manufacturer’s protocol. Subsequently, the EBOV libraries were subjected to twelve cycles of PCR amplification using universal Illumina primers that recognize the ligated adapter sequence of the libraries and enzymes from the Illumina TruSeq DNA Nano LP kit. PCR fragments were purified with Agencourt AMPure XP beads (Beckman Coulter) and eluted with Resuspension Buffer provided in the Illumina TruSeq DNA Nano LP kit. The DNA concentration of the library was determined using the dsDNA HS Assay Kit (Invitrogen) on a Qubit 4.0 Fluorometer (Invitrogen). The enriched libraries were sequenced without further size selection on a MiSeq instrument (Illumina) using the MiSeq Reagent Kit v3 (600-cycle) (Illumina). To balance GC content and low genomic diversity of the EBOV libraries, pools of up to five libraries were spiked into bacterial genomic MiSeq runs at 5%. For those brain samples from which reads were recovered, the average read depths ranged from 54 to 4,176, the median read depth ranged from 40 to 2,416, and the interquartile range varied from 32 to 3,444. For those liver samples from which reads were recovered, the average read depth ranged from 216 to 13,102, the median read depth ranged from 86 to 8,857, and the interquartile range varied from 216 to 10,498.

Raw sequence files were mapped using the nf-ViralMutations pipeline (https://github.com/phac-nml/nf-ViralMutations) v1.0.1 using KJ660347.2 as the reference sequence. Single nucleotide polymorphism (SNP) calling was manually verified and curated. The curated SNPs were imported into R v4.3.3 (49) and plotted using the tidyverse package v2.0.0 (50). Visuals were annotated in Inkscape v1.2.2.

### Transcriptomics

RNA extracted from the brains of eight ferrets from the second challenge experiment were subjected to RNA sequencing. RNA quality was assessed on a Bioanalyzer using the Agilent RNA 6000 Pico Kit (Agilent Technologies), and quantified using the RNA High Sensitivity Assay Kit (Invitrogen) on a Qubit Flex fluorometer (Invitrogen). Samples were evaluated for DNA contamination using the dsDNA HS Assay Kit (Invitrogen). An aliquot of diluted ERCC ExFold RNA Spike-Mix 1 (Invitrogen) was added to the RNA, per manufacturer’s guidelines, and the RNA was DNase-treated using ezDNase enzyme (Invitrogen). RNA-seq libraries were prepared from 100, 500, or 1000 ng of the DNase-treated RNA using the TruSeq Stranded mRNA Library Prep Kit (Illumina) following a slightly modified manufacturer’s protocol, where Superscript II was replaced with Superscript III reverse transcriptase to generate cDNA. The purified cDNA was then adenylated at the 3’ end and ligated to IDT for Illumina TruSeq RNA Unique Dual index adapters (Illumina). DNA fragments containing adapters on both ends were enriched via PCR and purified using AMPure XP beads (Beckman Coulter Genomics). Libraries were quantified on a Qubit Flex fluorometer using the dsDNA BR Assay Kit (Invitrogen), and DNA size distribution was assessed on a Bioanalyzer using the Agilent High Sensitivity DNA Kit (Agilent Technologies). Libraries were sequenced on a NextSeq 2000 (Illumina) using the NextSeq™ 2000 P3 Reagent Kit (100-cycle) (Illumina) for 1 x 100 bp reads. The reads were aligned to the ENSEMBL ferret genome using HiSAT.

Data analysis and visualization was performed using R v4.4.1 (49). Briefly, aligned data was normalized based on the ERCC spike-in control material using the BRGenomics v1.17.1 package (51) and then normalized further by regularized log transformation within the DESeq2 v1.46.0 package (52). The log_2_ fold change was centered for each gene, and the fifty genes with the highest variance were used to generate heatmaps using pheatmap v1.0.12 (53). For genes with no names, ENSEMBL identifiers were queried against the UniProt database (https://www.uniprot.org/) to determine if there were homologous genes found in other species. As some genes did not have annotated names, a heatmap displaying the top fifty named genes with the highest variance was also generated. Gene names were queried against the GOslim database using GO Term Mapper (https://go.princeton.edu/cgi-bin/GOTermMapper) to summarize the shared ontologies of the genes identified. Additional R packages used included dplyr v1.1.4 (54), rtracklayer v1.66.0 (55), and tidyverse v2.0.0 (50).

### Proteomics

Brain tissue homogenates were combined with an equal volume of 4X SDS Buffer (0.25 M Tris-HCl, pH 7.5, 4% SDS, 35% glycerol, 0.5% (w/v) bromophenol blue) and boiled for 10 minutes at 100°C prior to protein extraction using a modified SPEED method (56). The lysed brain homogenate (50 µL) was first acidified with 100 µL of trifluoroacetic acid (TFA) and after 15 minutes of shaking, was neutralized with 1,350 µL of a 20 mL solution of 2 M Tris base containing a cOmplete™ EDTA-free Protease Inhibitor Cocktail (PIC) tablet (Roche). Reduction and alkylation was performed separately with 10 mM dithiothreitol (DTT) and 55 mM iodoacetamide (IAA). Following an overnight acetone precipitation, samples were quantified by bicinchoninic acid (BCA) assay (Thermo Scientific).

Each sample (100 µg) was brought to 5% Sodium Dodecyl Sulfate (MP Biochemicals, Solon OH) in a total volume of 100 µl. The samples were processed using S-Trap mini spin columns (Protifi, Fairport NY USA) according to the manufacturer’s protocol. The samples were digested overnight with trypsin (Pierce, Rockford IL). The following morning, the peptides were concentrated to near-dryness (1-5 µl) using a vacuum centrifuge and labeled with the TMT 18-plex Isobaric Label Reagent Set (Thermo Fisher Scientific, Waltham, USA) per the manufacturer’s protocol; the TMT-labelled peptides were then mixed one-to-one. TMT sample mixtures were resuspended in 40 µL of buffer A (20 mM ammonium formate) and fractionated by high-pH C18 reversed-phased liquid chromatography using an Agilent 1260 Infinity liquid chromatography system (Agilent Technologies, Santa Clara, CA) equipped with a XBridge C18 3.5-µm, 2.1 x 100-mm column (Waters, Wexford Ireland) using a linear gradient of 20-56% buffer B (20 mM ammonium formate, 90% acetonitrile) over 52.5 minutes. The twelve TMT fractions were concentrated to near-dryness using vacuum centrifugation, and resuspended in 80 µL of nano buffer A (2% acetonitrile, 0.1% formic acid).

Each fractionated sample was analyzed by a liquid chromatography-tandem mass spectrometry using a nano-flow Easy-nLC 1200 connected in-line to an Orbitrap Fusion Lumos mass spectrometer (Thermo Fisher Scientific) with the ion source set to 2.1 kV. The samples were loaded on a 50-cm Easy-Spray c18 column (Thermo Fisher Scientific, ES903) with buffer A (2% acetonitrile, 0.1% formic acid) and run using a 120-minute gradient of 3% to 100% buffer B (80% acetonitrile, 0.1% formic acid). The following settings were used for data acquisition. The precursor scans were acquired in positive mode in the orbitrap with range of 375-1500 m/z at 120,000 resolution. Fragmentation scans were done in the IonTrap using CID fixed at a 35% collision energy with a 1.2 m/z isolation window, fragmenting as many targets as possible in a 2 second window. Each of the targets were also fragmented again using MS3 with HCD set to 65% normalized collision energy, 5 dependent scans, and analyzed in the orbitrap with a resolution set to 50,000.

Data was searched against the UniProt database for *Mustela putorius* (taxon ID: 9669) using Sequest within the Proteome Discoverer v3.1 software (Thermo Scientific). Subsequent data analysis and visualization was performed using R v4.4.1 (49). Briefly, data was normalized using variance stabilizing transformation within the DEP v1.28.0 package (57). Similarly to the transcriptomic data, the log_2_ fold change was then centered for each protein and the fifty proteins with the highest variance were used to generate heatmaps as above. Protein identifiers were queried using GO Term Mapper, as above, to summarize the shared ontologies of the proteins identified. Additional R packages used not listed above included readxl v1.4.3 (58) and stringr v1.5.1 (59).

### Histopathology, Immunohistochemistry (IHC), and *in situ* Hybridization (ISH)

Ferret brain samples were fixed in 10% neutral buffered formalin. Tissues were processed in a HistoCore Pearl Tissue Processor (Leica) followed by paraffin embedding with the Arcadia H/C embedding station (Leica). Sections were cut on a Leica model RM2255 microtome at 5 μm and left to dry overnight at 39°C.

To detect EBOV genomic RNA in formalin-fixed, paraffin-embedded tissues, RNA *in situ* hybridization (ISH) was performed using the RNAscope 2.5 HD RED kit (Advanced Cell Diagnostics, Newark, CA) according to the manufacturer’s instructions and USAMRIID SOP PT-04-10. Briefly, an ISH probe targeting the genomic fragment of EBOV nucleoprotein (NP) gene was designed and synthesized by Advanced Cell Diagnostics (Cat # 448581, Advanced Cell Diagnostics). Tissue sections were deparaffinized with xylene, underwent a series of ethanol washes and peroxidase blocking, and were then heated in kit-provided antigen retrieval buffer and then digested by kit-provided proteinase. Sections were exposed to ISH target probe and incubated at 40°C in a hybridization oven for 2 h. After rinsing, ISH signal was amplified using kit-provided pre-amplifier and amplifier conjugated to alkaline phosphatase and incubated with a Fast Red substrate solution for 10 minutes at room temperature. Sections were then stained with hematoxylin, air-dried, and mounted.

EBOV immunohistochemistry (IHC) was performed using the Dako Envision system (Dako Agilent Pathology Solutions, Carpinteria, CA, USA). After deparaffinization, rehydration, and methanol-hydrogen peroxide blocking, slides were stained using a mixture of USAMRIID mouse monoclonal anti-EBOV GP1,2 (M-DA01-A5-B11) antibody at a dilution 1:8,000 and USAMRIID mouse monoclonal anti-VP40 antibody (B-MD04-BD07-AE11) at a dilution 1:8,000, followed by a horseradish peroxidase-conjugated secondary anti-mouse polymer (Cat # K4003, Dako Agilent Pathology Solutions). All slides were exposed to brown chromogenic substrate, 3,3’-diaminobenzidine (DAB; K3468, Dako Agilent Pathology Solutions), counterstained with hematoxylin, dehydrated, and coverslipped. All images were collected using an Olympus BX46 microscope.

### Blood Biochemistry and Hematology

Complete blood cell counts in EDTA blood were performed using the VetScan HM5 hematology analyzer (Abaxis Veterinary Diagnostics) where lymphocytes, neutrophils, monocytes, white blood cells, and platelets were enumerated. Blood biochemistry was analyzed in lithium heparin blood use the VetScan VS2 chemistry analyzer (Abaxis Veterinary Diagnostics), and the following analytes were quantified: alanine aminotransferase (ALT), alkaline phosphatase (ALP), total bilirubin, albumin, globulin, glucose, total protein, amylase, blood urea nitrogen (BUN), potassium (K^+^), calcium (Ca^2+^), sodium (Na^+^), inorganic phosphate, and creatinine. For each parameter, the normal range was depicted as the mean ± standard deviation from all pre-infection samples.

### Quantification of Cytokines

Cytokines in plasma samples were quantified with the Ferret Cytokine Panel 1A kit (Cat # F103-Ka, Ampersand Biosciences) following the manufacturer’s instructions. Each sample was diluted 1:3 in sample diluent buffer and assayed in duplicate on a Luminex MAGPIX instrument. Data analysis was performed with the xPONENT software v4.2 (Luminex). The following cytokine levels were assessed: interferon gamma-induced protein 10 (IP-10; CXCL10), monocyte chemoattractant protein-1 (MCP-1; CCL2), macrophage inflammatory protein-1 (MIP-1; CCL4), interferon alpha (IFN-α), tumour necrosis factor alpha (TNF-α), interleukin 2 (IL-2), IL-4, IL-6, IL-8, IL-17, IL-12p40, and IL-12p70.

### ELISAs

EBOV GP-specific IgG levels in serum samples were quantified by indirect enzyme-linked immunosorbent assays (ELISA), as previously described (60). Briefly, half-area high-binding 96-well assay plates (Corning) were coated at 4°C overnight with 30 µl of 1 µg/ml of transmembrane domain-deleted EBOV GP protein (IBT Bioscience) prepared in carbonate buffer (pH 9.5). The plates were then blocked with 100 ul of 5% skim milk (BD Biosciences) prepared in PBS at 37°C for 1 h. Two-fold serial dilutions (starting at 1:200) of serum samples prepared in 2% skim milk were added to the plates for incubation at 37°C for 1 h, followed by washing with PBS containing 0.1% Tween-20 (PBST). The plates were incubated at 37°C for 1 h with an HRP-conjugated goat anti-ferret IgG secondary antibody (Novus) diluted 1:70,000 in 2% skim milk, and then washed again with PBST. Following addition of TMB solution (Life Technologies), chemiluminescence signal as measured at 650 nm using the Synergy HTX plate reader (Biotek) after 30 min. Endpoint dilution titers were calculated by determining the highest dilution that gave an average OD 650 reading greater than or equal to the cut-off OD value, which was set as the mean OD value for pre-infection serum samples plus three times the standard deviation.

The levels of human IgG (i.e., 1C3 and 1C11) were quantified in serum samples from the two ferret challenge experiments as well as the antibody pharmacokinetics experiment using the Human IgG ELISA Kit (Cat # ab195215, Abcam), according to manufacturer’s instructions. A standard curve was generated in GraphPad Prism and used to calculate the concentration of total human IgG in each sample. For the pharmacokinetics experiment, the half-life of each antibody in each animal was calculated using a one-phase decay curve incorporating data from the 96-504 hour time points. Because human antibody was not detected at any time point in one of the animals treated with 1C11, the half-life for this mAb was calculated from only three animals.

### Statistical Analyses

GraphPad Prism (version 10) was used to perform all statistical tests (as indicated in the figure legends) and generate all graphs, with the exception of the transcriptomic and proteomic analyses, which used R v4.4.1 (49). Statistically significant differences are indicated with asterisks, where a *P* value less than or equal to 0.05 is marked with one asterisk (*), less than or equal to 0.01 marked with two asterisks (**), less than or equal to 0.001 marked with three asterisks (***), and less than or equal to 0.0001 marked with four asterisks (****).

## ACKNOWLEDGEMENTS

We thank Zalgen Labs for producing the monoclonal antibodies used in this study. We thank the Veterinary Technical Services team at the National Microbiology Laboratory (NML) for animal care and technical assistance, and Stuart McCorrister and Dr Christopher Grant at the NML for the proteomics/mass spectrometry methods and data acquisition. We also thank the Genomics Core Facility at the NML for technical assistance.

This work was supported by the Public Health Agency of Canada (PHAC) as well as a grant from the National Institute of Allergy and Infectious Diseases (NIAID), U19 AI142790-03, Consortium for Immunotherapeutics against Emerging Viral Threats (L.B., E.O.S.). This work was also supported by the Joint Science and Technology Office for Chemical and Biological Defense (JSTO-CBD) of the Defense Threat Reduction Agency (DTRA; CB11408 to X.Z.).

## AUTHOR CONTRIBUTIONS

**Conceptualization**: Wenguang Cao, Shihua He, Logan Banadyga

**Methodology**: Wenguang Cao, Shihua He, Helene Schulz, Jordan Wight, Karla Emeterio, Jonathan Audet, Kathy Frost, Lilianne Gee, Peter McQueen, Patrick Chong, Garrett Westmacott, Xiankun Zeng, Logan Banadyga

**Investigation**: Wenguang Cao, Shihua He, Helene Schulz, Jordan Wight, Michael Chan, Karla Emeterio, Guodong Liu, Jonathan Audet, Kevin Tierney, Kimberly Azaransky, Kathy Frost, Lilianne Gee, Peter McQueen, Patrick Chong, Garrett Westmacott, Xiankun Zeng, Logan Banadyga

**Formal analysis**: Wenguang Cao, Helene Schulz, Jordan Wight, Karla Emeterio, Jonathan Audet, Lilianne Gee, Peter McQueen, Patrick Chong, Garrett Westmacott, Xiankun Zeng, Logan Banadyga

**Writing – original draft**: Wenguang Cao, Jordan Wight, Jonathan Audet, Xiankun Zeng, Logan Banadyga

**Writing – review and editing**: All authors

**Visualization**: Wenguang Cao, Helene Schulz, Jordan Wight, Jonathan Audet, Xiankun Zeng, Logan Banadyga

**Supervision**: Stephanie Booth, Garrett Westmacott, Logan Banadyga

**Resources**: Stephanie Booth, Garrett Westmacott, Erica Ollmann Saphire, Xiankun Zeng, Logan Banadyga

**Funding acquisition**: Erica Ollmann Saphire, Xiankun Zeng, Logan Banadyga

## SUPPORTING INFORMATION CAPTIONS

**S1 Figure. Schematic of the ferret treatment experiments.** mAb efficacy evaluation was performed over two independent experiments, with the “early” and “late” antibody treatments tested in one experiment (A, B) and the “very early” and “mid” treatments tested in a second experiment (C, D). All animals were inoculated on 0 days post-infection (DPI) with a target dose of 1000 TCID_50_ EBOV via either the IN or the IM route, after which they were treated with 30 mg/kg each of antibody on 2 and 5 DPI (“early”) (A), 3 and 6 DPI (“late”) (B), 2 and 4 DPI (“very early”) (C), or 3 and 5 DPI (“mid”) (D). Blood and swab samples were collected from all animals on all days indicated on the schematic. The first experiment ended on 29 DPI (A, B), while the second experiment ended on 36 DPI (C, D).

**S2 Figure. Virus genome sequences from acute and atypical ferrets.** Viral RNA was isolated from brain (A) and liver (B) tissue samples for each of the indicated animals and subjected to next generation sequencing. The frequency of each identified mutation (compared to the EBOV/Mak-C07 reference sequence, Gen Bank Accession No. KJ660347.2) is indicated by a coloured rectangle on the heatmap, with animal ID on the y-axis and genomic mutations on the x-axis. Nucleotide changes are defined across the bottom of each heatmap, using their position in the genome and lowercase lettering for the mutated base pairs. Corresponding amino acid changes are defined across the top of each heatmap, using the amino acid number and the uppercase, single-letter codes for the amino acids. Grey rectangles indicate that no sequencing data was obtained at this position; crossed lines indicate no difference compared to the reference sequence. Sequence data from the stock virus, which was used to inoculate the ferrets, is provided for comparison.

**S3 Figure. Transcriptomic analyses of ferret brain tissue for the fifty most variable expressed genes.** Brain tissue samples from a subset of ferrets (all from the second challenge experiment) were subjected to transcriptomic analysis. A heatmap shows the differential expression of the fifty genes with the most variable expression, including those without names or known functions, shown as log_2_ fold change centered for each gene.

**S4 Figure. Additional blood biochemistry parameters for acute, atypical, and surviving ferrets.** Levels of amylase (A, B), blood urea nitrogen (BUN) (C,D), K^+^ (E, F), Ca^2+^ (G, H), Na^+^ (I, J), inorganic phosphate (K, L), and creatinine (M, N) were quantified in all blood samples from all animals. Levels of each of these analytes are depicted over time (A, C, E, G, I, K, M). The vertical area shaded grey, from 12 to 18 DPI, represents the window in which atypical disease was observed, while the horizontal area shaded grey represents the normal range for each parameter. The levels of each analyte at the terminal time points are also depicted separately (B, D, F, H, J, L, N), with means and standard deviations indicated. Mean levels were compared using a one-way ANOVA with Tukey’s multiple comparison test. ns, not significant; *, p≤0.05; **, p≤0.01; ***, p≤0.001; ****, p≤0.0001.

**S5 Figure. Additional cytokine levels in acute, atypical, and surviving ferrets.** Levels of IFN-α (A, B), TNF-α (C, D), IL-12p70 (E, F), IL-17 (G, H), and IL-2 (I, J) were quantified in all serum samples from all animals via Luminex assay. Levels of each cytokine are depicted over time (A, C, E, G, I). The vertical area shaded grey, from 12 to 18 DPI, represents the window in which atypical disease was observed. The levels of each cytokine at the terminal time points are also depicted separately (B, D, F, H J), with means and standard deviations indicated. Mean levels were compared using a one-way ANOVA with Tukey’s multiple comparison test. ns, not significant; *, p≤0.05.

**S6 Figure. Pharmacokinetics of 1C3 and 1C11 in ferrets.** Ferrets were injected intraperitoneally with 30 mg/kg of either 1C3 (n = 4) or 1C11 (n = 4), after which serum samples were collected at 1, 6, 24, 48, 72, 96, 120, 144, 168, 336, and 504 hours post-treatment. Total human IgG (i.e., 1C3 or 1C11) concentrations were quantified in each serum sample via ELISA and depicted over time for each animal (A). Note that human antibodies could not be detected at any time point in one animal injected with 1C11; for this reason, data from this animal were excluded from the analysis. Antibody half-lives were calculated for each animal injected with 1C3 (B) or 1C11 (C) based on the data from 96 to 504 hours using a one-phase decay model to better capture the slow elimination phase. The animal ID is provided for each decay curve, with the sex of the animal indicated (M, male; F, female).

**S1 Table. Summary of Experiments #1 and #2**

